# p16^High^ senescence restricts totipotent potential during somatic cell reprogramming

**DOI:** 10.1101/2022.08.24.504108

**Authors:** Bogdan B. Grigorash, Dominic van Essen, Laurent Grosse, Alexander Emelyanov, Benoît Kanzler, Clement Molina, Elsa Lopez, Oleg N. Demidov, Carmen Garrido, Simona Saccani, Dmitry V. Bulavin

**Affiliations:** Institute for Research on Cancer and Aging of Nice (IRCAN); INSERM; Université Côte d’Azur, CNRS, Nice, France; Max Planck Institute of Immunobiology and Epigenetics, Stübeweg 51, D-79108 Freiburg, Germany; INSERM UMR1231, LopSTIC, University of Burgundy Franche-Comté, Dijon, France

## Abstract

The discovery of four factor (4F)-induced reprogramming of somatic cells into induced pluripotent stem (iPS) cells has revolutionized the fields of cell and regenerative biology. In contrast, the feasibility of a direct conversion of somatic cells into a totipotent state defined as the ability to produce all cell types of an organism, including extraembryonic tissues, is not well established. Using genetic and chemical approaches to manipulate senescent cells, here we found that removal of p16^High^ cells resulted in 4F-induced reprogramming of somatic cells into totipotent-like stem cells. These cells expressed markers of both pluripotency and the 2-cell (2C) embryonic state, readily formed implantation-competent blastocyst-like structures, blastoids, and following morula aggregation, contributed to embryonic and extraembryonic lineages in E12.5 embryos. We identified senescence-dependent regulation of nicotinamide N-methyltransferase (NNMT) as a key mechanism controlling the S-adenosyl-l-methionine (SAM) levels during 4F-induced reprogramming that was required for expression of the 2C genes and acquisition of an extraembryonic potential. Our results show that the presence of p16^High^ senescent cells, high NNMT and low SAM limit cell plasticity during 4F-reprogramming, while their modulation could help to achieve the highest state of stem cell potency, totipotency.

## Introduction

Research into rejuvenation and life extension with the aim of developing relevant therapies is undergoing a renaissance. Several recent discoveries have demonstrated that the fitness of aged tissues can be noticeably improved by inducing their molecular and epigenetic rejuvenation *in vivo*. One such approach has been to leverage the 4 Yamanaka reprogramming factors (4F) to counteract the *in vivo* signs of aging and increase the life span of mice with a premature aging disease(Hishida et al., 2022)(Reddy et al., 2021)(Ocampo et al., 2016). The concern however is that epigenetic reprogramming and its impact on rejuvenation could be significantly attenuated or even lost in fully aged tissues due to profound changes in the epigenome of old cells, persistent chronic inflammation, significant tissue fibrosis and other age-related debilitating conditions. In this respect, recent findings highlight the important role of a continuous build-up of senescent cells as one of the key drivers of aging-induced tissue deterioration, while removal of some senescent cells showed great promise in ameliorating several age-related conditions and extending life span in mice (Baker et al., 2011)(Baker et al., 2016)(Childs et al., 2017)(Xu et al., 2018). How 4F reprogramming and senescence interconnect *in vivo*, however, remains largely under-investigated.

Initial observations revealed a negative role for cellular senescence in somatic cell reprogramming *in vitro*(Li et al., 2009)(Banito et al., 2009)(Utikal et al., 2009). More recent studies, however, using genetic backgrounds to attenuate senescence showed that while p53 deficiency recapitulated the negative role of senescence in reprogramming, an unexpected reduction in the efficiency of *in vivo* teratoma formation was found in Cdkn2a-deficient mice(Mosteiro et al., 2016). Deletion of either Cdkn2a/p16 or p53 gene both have a strong impact on senescence in multiple cell types, and thus the exact reason for differences in the *in vivo* reprogramming between these two genetic backgrounds is unclear. One explanation is that Cdkn2a deficiency has an impact on macrophage biology including enhanced antiinflammatory M2 polarization (Cudejko et al., 2011). In turn, teratomas developed on a Cdkn2a deficient background contain far fewer macrophages which is not the case for p53-deficient mice (Mosteiro et al., 2016) while macrophage depletion significantly attenuate teratoma development, indicating their positive contribution to teratoma growth (Chen et al., 2014). Whether other approaches to attenuate senescence including removal of p16^High^ cells have a similar to Cdkn2a deficiency suppressive effect on the *in vivo* iPS reprogramming remains unclear.

Here, we employed genetic and chemical approaches to selectively deplete p16^High^ senescent cells and discovered their critical role in iPS reprogramming *in vivo* as well in achieving a totipotent-like state. In addition to upregulation of several cardinal 2C factors, including Dux and MERVL (Morgani et al., 2013)(Genet and Torres-Padilla, 2020), totipotent-like cells described here expressed Line-1 retro-elements and Stella/Dppa3, further re-enforcing their totipotent state. Mechanistically, we identified senescence-dependent metabolic-epigenetic reprogramming through the NNMT-dependent regulation of SAM levels, expression of the 2C genes and global histone methylation. Here, we found a critical role for p16^High^ senescent cells, NNMT and SAM in reprogramming into a totipotent state, a mechanism that could be applicable to achieve a higher cell plasticity under different conditions.

## Results

### Expression of the four reprogramming factors is accompanied by accumulation of p16^High^ senescent cells

Cell passaging *in vitro* results in activation of the p16 *(Cdkn2a)* gene and emergence of SA-ß-gal positive cells defining replicative senescence. To assess the role of iPS reprogramming in the induction of senescence, we isolated primary dermal fibroblasts (DF) from i4F;rtTA mice and cultured them with and without doxycycline (DOX) until the iPS colonies were formed(Abad et al., 2013). The DOX-inducible system allowed us to control the expression of four reprogramming factors while avoiding any potential impact of retroviral infection on the activation of p16. On day 15, approximately 10% of DOX-treated DF were positive for SA-ß-gal compared to only 1% of non-treated DF, while p16 mRNA expression was increased by >20-fold (Figure 1A).

**Figure 1.**
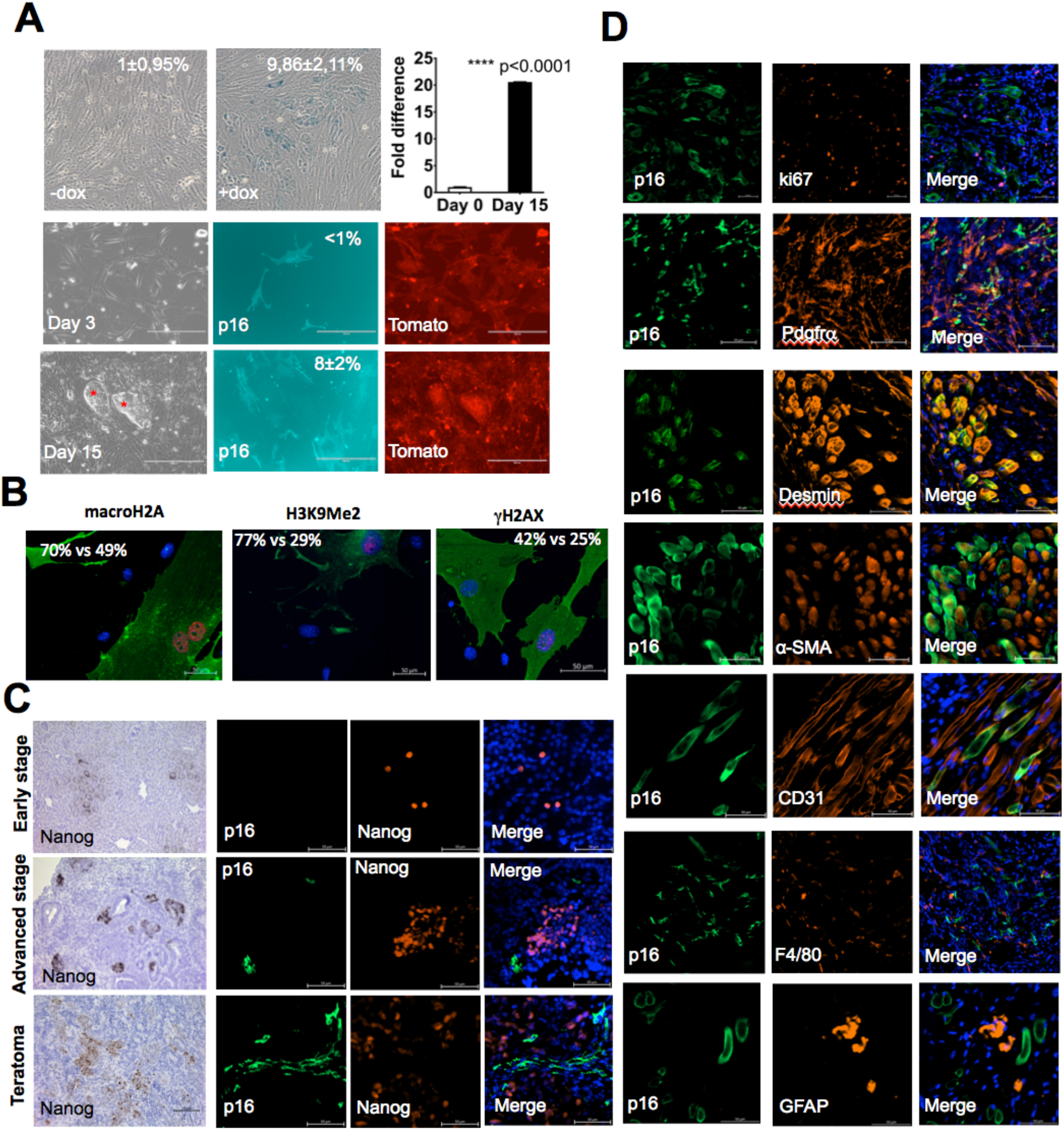
p16^High^ cells have a negative impact on iPS reprogramming. **A** (top left) – analysis of SA-β-galactosidase (SA-β-gal+ cells ± SD) activity on d15 of DF untreated (left) and treated (right) with DOX. Scale bar - 50 μm. Top right–qPCR analysis of p16 mRNA levels (± SD) in DFs on d15 of DOX treatment. P value estimated by unpaired, two-tailed Student’s t-test with Welch’s correction. Middle panel and bottom panel - p16Cre;i4F;rtTA;mTmG DF were treated with DOX, p16^High^ cells (mean ± SD) were counted on d3 and d15. iPS colonies marked with red asterisks. Scale bar – 200 μm. **B** – Representative images of co-staining of p16^High^ cells on d15 of reprogramming with macroH2A, H3K9Me2 and γH2AX, markers of senescence. Numbers represent a fraction of cells expressing high level of senescence markers in p16^High^ versus p16^Low^ cells. Scale bar – 50 μm. **C** (left panels) – representative images of mouse kidney at different stages of teratoma formation stained for Nanog. Scale bar - 100 μm. Right panels - same reprogramming stages co-stained for Nanog and EGFP-p16^High^. Scale bar - 50 μm. **D** - immunofluorescent analysis of teratomas for p16^High^ cells and proliferative marker Ki67, mesenchymal markers Pdgfrα, Desmin and α-SMA, vascular endothelial marker Cd31, macrophage marker F4/80 and neuroectodermal marker Gfap. Scale bar – 50 μm.

Next, we crossed p16Cre;mTmG (Grosse et al., 2020) and i4F;rtTA (Abad et al., 2013) mice and established DOX-reprogrammable p16^High^ reporter DF lines. The p16Cre;mTmG reporter system allowed us to perform lineage tracing of p16^High^ cells and to analyze EGFP^+^/p16^High^ cells in greater detail. As shown in Figure 1A, EGFP^+^/p16^High^ cells accounted for 8±2% of the population on d15 of iPS reprogramming. Lineage tracing of p16^High^ cells revealed that p16 activation was incompatible with iPS reprogramming as 100% of the iPS colonies arose from the Tomato^+^/EGFP^-^ population (Figure 1A, shown with asterisks), implying that the level of p16 in reprogrammed cells was insufficient to induce mTmG recombination. Analysis of different senescence markers on d12 of reprogramming (prior to the formation of any visible iPS clones) revealed an increase in the number of γH2Ax^+^ foci, K9me2 and macroH2a nuclear staining in EGFP-positive/p16^High^ compared with EGFP-negative/p16^Low^ cells (Figure 1B). Once the iPS cultures are established, we observed no EGFP^+^/p16^High^ cells appearance after multiple passages implying that in *in vitro* reprogramming experiments p16^High^ senescent cells are only present during the establishment of iPS colonies and not their maintenance.

### p16^High^ senescent cells appear later than Nanog^+^ cells in vivo and are of mesenchymal origin

Next, we investigated iPS reprogramming using a genetic approach to label p16^High^ cells *in vivo*. The reprogramming process was induced *in vivo* by the addition of DOX to the drinking water of p16Cre;i4F;rtTA;mTmG mice and the rate of EGFP^+^/p16^High^ cell appearance was then assessed at different stages. Since the pancreas and the kidney are the main sites of teratoma formation during *in vivo* reprogramming (Abad et al., 2013), we analyzed both organs from p16Cre;i4F;rtTA;mTmG mice at different stages of teratoma development after 3 weeks of treatment with DOX. Immunohistochemical analysis of Nanog^+^ cells at early stages of reprogramming (Figure 1C, top left panel) revealed that in spite of presence of Nanog^+^ cells, the typical morphology of kidney tissue was largely preserved and there were no major pathological changes associated with teratoma growth. At this stage, EGFP^+^/p16^High^ cells were completely absent in the kidney (Figure 1C, top right panels). Next, we identified kidneys with the onset of teratoma growth, which we regarded as an advanced stage typically containing one or two germ layer derivatives. At this stage, our analysis (Figure 1C, middle panels) showed loss of the typical kidney tissue morphology with a large number of Nanog^+^ cells, predominantly organized in clusters. EGFP^+^/p16^High^ cells appeared for the first time at this stage. Further analysis of fully developed teratomas showed a change in the pattern of Nanog^+^ cells from small clusters to large uniform distributions and a larger overall number of EGFP^+^/p16^High^ cells (Figure 1C, low panels). H&E staining revealed the presence of derivatives of all three germ layers in teratomas at this stage (SFigure 1A), which further indicated that these were derived from fully reprogrammed pluripotent cells. Thus, we showed that Nanog^+^ cells appear before p16^High^ cells *in vivo*.

We confirmed the mitotically inactive status of EGFP^+^/p16^High^ cells by the lack of costaining with Ki67, a prominent marker of cell proliferation outside of the G_0_ phase of the cell cycle (Figure 1D, top panel). This provided further evidence that EGFP^+^/p16^High^ cells in reprogrammed teratomas are non-dividing, which is a feature of senescent cells. Next, we analyzed the cellular origin of EGFP^+^/p16^High^ cells in the teratoma. We speculated that these cells could have activated p16 in response to hyperproliferation during teratoma growth due to signals from the microenvironment, or were derived from reprogrammed cells including iPS, or both. Therefore, we studied teratoma developed during *in vivo* reprogramming of p16Cre;i4F;rtTA;mTmG mice. Macrophages express high levels of p16 under certain conditions and their migration to the teratoma site is critical for teratoma growth(Hall et al., 2017)(Grosse et al., 2020)(Chen et al., 2014). To check the possibility that EGFP^+^/p16^High^ cells are macrophages, we performed co-staining of the macrophage marker F4/80. To check the possibility that EGFP^+^/p16^High^ cells are endothelial cells of blood vessels (Grosse et al., 2020), we used an anti-CD31 antibody. In addition, we carried out a co-staining analysis for the neuroectodermal marker Gfap. Finally, we performed immunostaining for several fibroblast-specific antigens (αSMA, Pdgfrα, Desmin). We found that only a very small fraction of the EGFP^+^/p16^High^ cell population was positive for F4/80 and CD31, while no co-staining with Gfap was detected (Figure 1D). In contrast, we observed a strong co-localization of EGFP^+^/p16^High^ cells with different fibroblast-specific markers (Figure 1D). Our data suggest that the majority of EGFP^+^/p16^High^ cells in the growing teratoma are of mesenchymal origin and specifically, are fibroblasts.

To determine whether EGFP^+^/p16^High^ fibroblasts are derived from the teratoma microenvironment or from reprogrammed cells including iPS, we then established cultures of p16Cre;i4F;rtTA;mTmG iPS cells under feeder-free culture conditions to avoid fibroblast contamination. We confirmed that 100% of the iPS clones established from p16 Cre;i4F;rtTA;mTmG fibroblasts were EGFP^-^/p16^Low^. Subsequently, we engrafted EGFP^-^/p16^Low^ iPS cells into NSG mice. In this model, the absence of EGFP^+^/p16^High^ cells in teratomas developed in NSG mice would confirm that EGFP^+^/p16^High^ cells are derived from the microenvironment. In contrast, we found numerous EGFP^+^/p16^High^ cells (SFigure 1B) comparable to the number of EGFP^+^/p16^High^ cells grown during *in vivo* reprogramming of p16Cre;i4F;rtTA;mTmG mice. Further co-staining revealed a similar mesenchymal origin of EGFP^+^/p16^High^ cells (SFigure 1B). Thus our analysis confirmed that the majority of EGFP^+^/p16^High^ cells originate from cells that undergone the reprogramming process specifically from iPS cells at the stage of teratoma formation *in vivo*. This was consistent with our finding that the first EGFP^+^/p16^High^ cells emerge only after the appearance of Nanog^+^ cells during *in vivo* reprogramming (Figure 1C).

### Genetic removal of p16^High^ senescent cells increases the in vitro iPS cell reprogramming efficiency

To further assess the consequences of p16 activation during iPS cell reprogramming, we used a genetic approach for selective elimination of p16^High^ cells. We crossed p16Cre;i4F;rtTA mice with a mouse strain containing a floxed STOP cassette upstream of the diphtheria toxin subunit A (DTA) gene. Once a DTA-expressing cell is killed and DTA is released into the environment, it is no longer toxic; therefore, this model facilitated targeted removal of p16^High^ cells without any negative influence on the surrounding p16^Low^ cells. First, we evaluated the accumulation of SA-ß-gal positive cells and p16 activation during reprogramming in DF isolated from p16Cre;i4F;rtTA;DTA mice (DTA DF). As shown in Figure 2A, selective removal of p16^High^ cells significantly reduced (from 10% to 3%) the proportion of SA-ß-gal positive cells on d15 of reprogramming. qPCR analysis further indicated a reduced level of p16 mRNA in DTA DF when compared to control DF (Figure 2A). Subsequent comparison of FACS-sorted EGFP^+^/p16^High^ and EGFP^-^/p16^Low^ DFs with DTA DFs showed that EGFP^+^/p16^High^ cells in reprograming cultures contained upregulated expression of p16 mRNA and multiple senescence-associated secretory phenotype (SASP) factors (Figure 2B). The mRNA levels of these factors were significantly reduced in DTA DFs, further supporting the efficiency of our genetic approach to the removal of p16^High^ senescent cells during iPS cell reprogramming.

**Figure 2.**
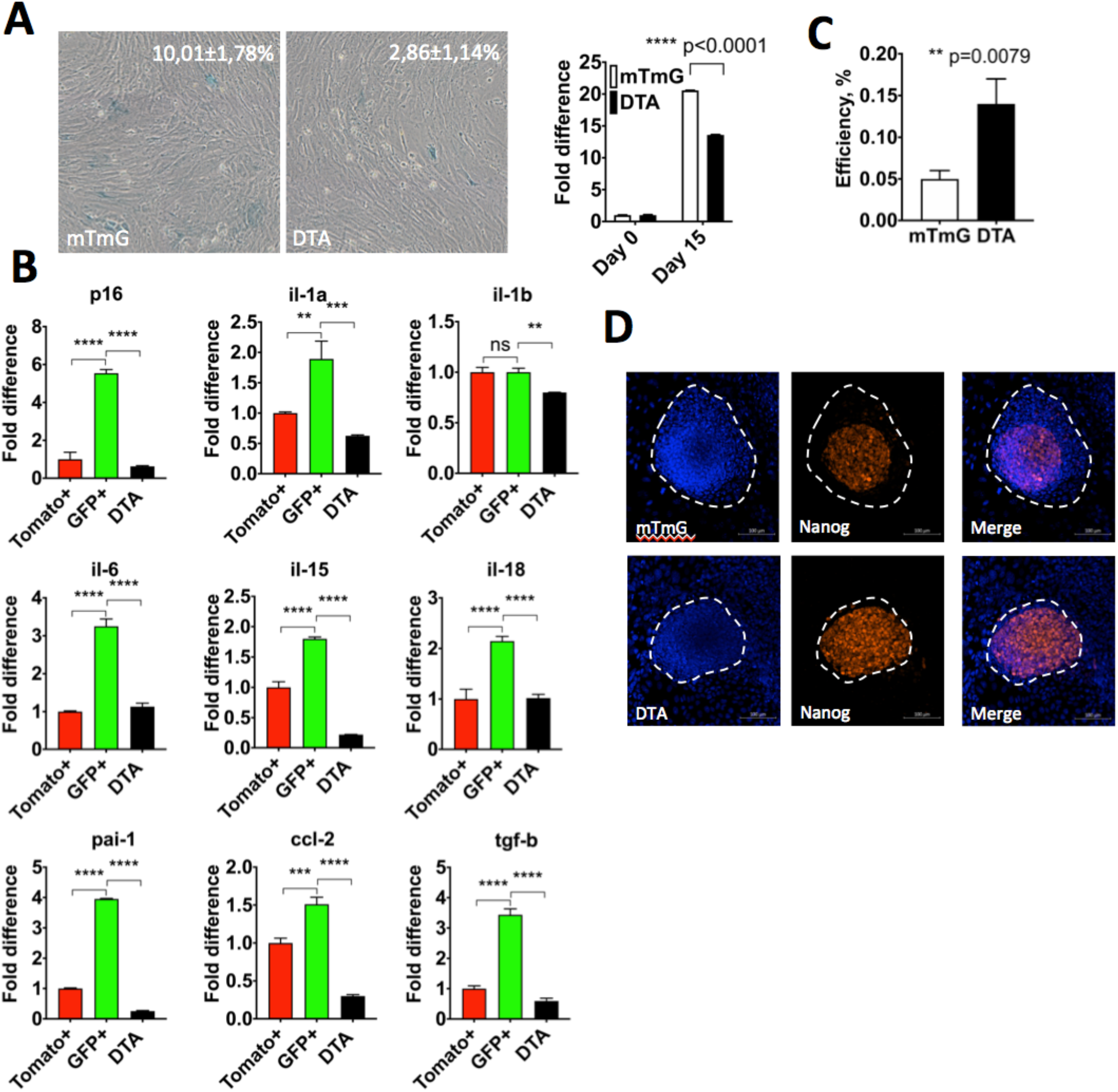
Genetic ablation of p16^High^ cells increases efficiency of iPS reprogramming *in vitro*. **A** (left panel) - analysis of SA-β-gal+ activity in DF without (mTmG) and with (DTA) ablation of p16^High^ cells (SA-β-gal+ cells ± SD) on d15 of DOX treatment. Right panel – qPCR analysis of p16 mRNA expression (± SD) in mTmG and DTA DF on d15. p-value estimated by unpaired, two-tailed Student’s test with Welch’s correction. **B** – qPCR analysis of p16 and SASP-related genes (± SD) in FACS-sorted GFP+ (p16^High^) and Tomato+ (p16^Low^) from p16Cre;i4F;rtTA;mTmG DFs and p16Cre;i4F;rtTA;DTA (DTA) DFs on d12. P-values estimated by ordinary one-way ANOVA with Sidak’s multiple comparisons test. ** p<0.005, *** p<0.0005, *** p<0.0001. **C** - iPS reprogramming efficiency in control DF (mTmG) and DTA DFs (± SD). p-value estimated by unpaired, two-tailed Student’s t-test. **D** – analysis of Nanog+ cells distribution within mTmG and DTA iPS colonies. Scale bar – 100 μm.

p16^High^ cells did not give rise to iPS colonies (Figure 1A), but appeared among surrounding cells with the potential to influence reprogramming via a paracrine signaling mechanism. To assess the contribution of p16^High^ cells in iPS cell reprogramming, we reprogrammed DF with and without a DTA cassette and counted the number of colonies positive for the ES cell marker, alkaline phosphatase. We found that DTA DF underwent reprogramming significantly more efficiently than the control DF (Figure 2C). Importantly, removal of p16^High^ senescent cells also significantly improved the quality of iPS clone reprogramming based on the number and distribution of Nanog+ cells, which appeared to be more homogenous throughout entire colony (Figure 2D).

Previously, we found that most EGFP^+^/p16^High^ cells detected in teratomas during *in vivo* reprogramming express markers of the mesenchymal lineage (Figure 1D, SFigure 1B). To assess the role of these cells in reprogramming *in vivo*, we isolated Pdgfrα-positive and -negative cells from teratomas developed in control (p16Cre;i4F;rtTA;mTmG) mice by magnetic-activated cell sorting and found that the majority of SA-ß-gal-positive cells were present in the Pdgfrα-positive fraction (Figure 3A). In turn, Pdgfrα-positive cells from DTA teratoma contained significantly fewer SA-ß-gal-positive cells compared to the number of Pdgfrα-positive cells from control teratomas. These data support the mesenchymal origin of EGFP^+^/p16^High^ cells and confirm the efficiency of senescent cell removal by our genetic DTA approach *in vivo*.

**Figure 3.**
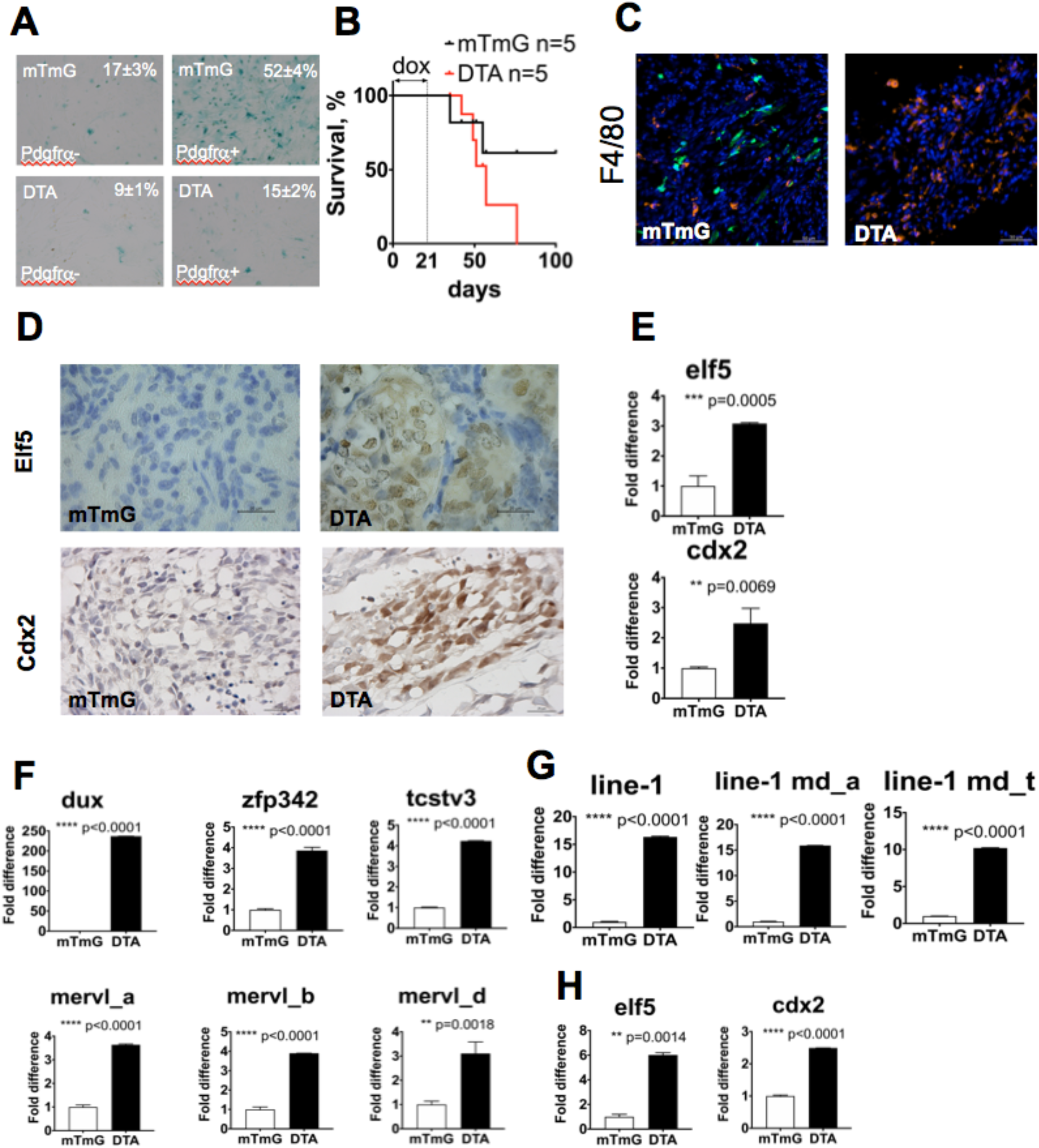
Removal of p16^High^ cells during *in vivo* and *in vitro* iPS reprogramming leads to emergence of cells with totipotent features. **A** - Staining for SA-β-gal (SA-β-gal+ cells ± SD) of FACS-sorted Pdgfrα+ and Pdgfrα– populations, isolated from mTmG and DTA teratomas. **B** - survival curve of p16Cre;i4F;rtTA;mTmG and p16Cre;i4F;rtTA;DTA mice, administered with DOX for 21d. **C** – analysis of macrophages (F4/80, Alexa 647) infiltration in mTmG and DTA teratoma. Scale bar represents 50 μm. GFP-p16^High^ cells. **D** - immunohistochemical analysis of trophectodermal markers Elf5 and Cdx2 in control (mTmG) and p16^High^-depleted (DTA) teratoma developed in 2-month-old mice. Scale bars: 20 μm. **E** - qPCR analysis of elf5 and cdx2 expression (± SD) in mTmG and DTA teratomas (2-month-old). p value calculated by unpaired, two-tailed Student’s t-test with Welch’s correction. **F** (top panel) - qPCR analysis of 2C genes in mTmG and DTA teratomas. Bottom panel – qPCR analysis of Mervl LTR retroelement, p-value evaluated by unpaired, two-tailed Student’s t-test with Welch’s correction. **G** – qPCR analysis of non-LTR retroelement line-1 and its 2 subclasses md_a and md_t. Statisctical analysis and data presentation same as in **F**. **H** - Expression of trophoblast stem cell-specific genes on d5 of trophoblast differentiation of mTmG and DTA iPS, p-value evaluated by unpaired, two-tailed Student’s t-test with Welh’s correction.

Since we observed more complete and efficient reprogramming of DF after genetic ablation of p16^High^ cells *in vitro*, we next compared the rate of teratoma growth in senescence-proficient (mTmG) and senescence-deficient (DTA) mice *in vivo* by administering DOX in the drinking water of p16Cre;i4F;rtTA;mTmG and p16Cre;i4F;rtTA;DTA mice respectively. Analysis of mouse survival recorded to the point at which all DTA mice were sacrificed because of teratoma formation revealed that >50% of mTmG mice remained teratoma-free (Figure 3B). Further analysis revealed the presence of macrophages in DTA teratomas in numbers that were comparable to those in teratomas developed in control mice (Figure 3C). Thus, removal of p16^High^ cells accelerates iPS cell reprogramming *in vitro* and teratoma development *in vivo* without affecting the recruitment of macrophages, which was in contrast to previous studies on a Cdkn2a-deficient background(Mosteiro et al., 2016).

### Genetic removal of p16^High^ cells during reprogramming results in expression of markers of extraembryonic tissues and the 2C embryonic stage

Next, we analyzed how removal of p16^High^ cells affects the pluripotent status of cells. H&E staining of teratomas developed in p16Cre;i4F;rtTA;DTA mice confirmed the presence of tissues representing all three germ cell layers (SFigure 2). We then extended our analysis to evaluate trophectodermal lineage cells in teratomas obtained from control and DTA mice. Trophectoderm formation is the first lineage decision during the pre-implantation development of the mouse embryo and pluripotent stem cells (ES or iPS cells) have insufficient “stemness” to differentiate into this lineage. To investigate this further, we stained cells for Elf5 and Cdx2, markers of trophectoderm lineage and transcription factors involved in trophoblast stem cell maintenance. We found numerous cells positive for both Elf5 and Cdx2 specifically in teratomas obtained from DTA but not mTmG mice (Figure 3D) and these results were confirmed by qPCR analysis (Figure 3E).

One possible explanation for the trophectodermal gene expression in teratomas developed in DTA mice is that in the absence of senescence, the stem cell potential of some, or even all iPS cells, is extended to a totipotent 2C-like state. Such cells in a 2C-stage embryo possess a number of unique properties that distinguish them from pluripotent stem cells including the activation of specific factors like Dux, Tcstv1, Tcstv3, as well as enhanced transcription of Mervl LTR retro-elements(Genet and Torres-Padilla, 2020). Thus, based on our preliminary analysis of Elf5 and Cdx2 expression in teratomas, we hypothesized that removal of senescent cells during reprogramming affects the potency of iPS cells making them competent to differentiate into any embryonic lineage; that is, these cells approach totipotency or, more precisely, exist in a state that closely resembles the 2C-stage (2C-like cells). Our qPCR analysis of numerous genes related to early mouse embryonic development in DTA teratomas indeed confirmed the activation Mervl retro-elements, as well as transcription factors implicated in the maintenance of 2C-like cells (Figure 3F). To our surprise, however, we also found a significant upregulation of non-LTR retro-elements, Line1, in DTA teratomas (Figure 3G).

To evaluate the ability of DTA iPS cells to develop similar extended 2C-like properties *in vitro, we* established feeder-free p16Cre;i4F;rtTA;mTmG (mTmG) and p16Cre;i4F;rtTA;DTA (DTA) iPS cell lines and induced their differentiation into trophoblast stem cells. We found that DTA iPS cells upon differentiation showed a significant upregulation of Elf5, and Cdx2 expression, confirming their ability to contribute to additional extraembryonic cell type *in vitro* (Figure 3H). Thus, establishment of iPS cells under senescence-free conditions results in the appearance of totipotent-like features both *in vitro* and *in vivo*.

### Senolytics dasatinib and quercetin prevent accumulation of p16^High^ cells during reprogramming and increases the iPS reprogramming efficiency

Next, we sought to confirm our results obtained by genetic ablation of p16^High^ senescent cells using a second approach. For that we decided to use a combination of dasatinib and quercetin (D+Q) as well as navitoclax, which are recently discovered drugs to selectively kill senescent cells, senolytics (Zhu et al., 2015)(Zhu et al., 2016). Of note, our analysis revealed that the published doses of navitoclax (>1 μM) (Zhu et al., 2016) were highly toxic during the first 3 days of iPS reprogramming when no senescent cells were present, requiring a significant reduction in concentration (<0.1μM). Thus, caution is necessary to avoid an anti-proliferative activity as a confounding factor in evaluation of the role of navitoclax-sensitive senescent cells in iPS reprogramming. As shown in Figure 4A, both D+Q and reduced doses of navitoclax prevented accumulation of SA-ß-gal positive cells during iPS reprogramming, while only D+Q reduced the number of EGFP^+^/p16^High^ cells (Figure 4B, left panel). Expression of p16 mRNA on d15 in cells treated with D+Q was also significantly reduced (Figure 4B, middle panel) and further analysis of the reprogramming efficiency revealed that cells treated with D+Q, but not with navitoclax, had more iPS colonies (Figure 4B, right panel). Taking into consideration the similar effects of D+Q to DTA in reprogramming (fewer SA-ß-gal^+^ cells, decreased p16 mRNA levels, more efficient reprogramming), we analyzed the distribution of the Nanog protein in iPS colonies. In accordance with our DTA experiments (Figure 2D), D+Q treatment resulted in a more even distribution of Nanog+ cells within the iPS cell colony, which is a feature of a better quality of reprogramming (Figure 4C).

**Figure 4.**
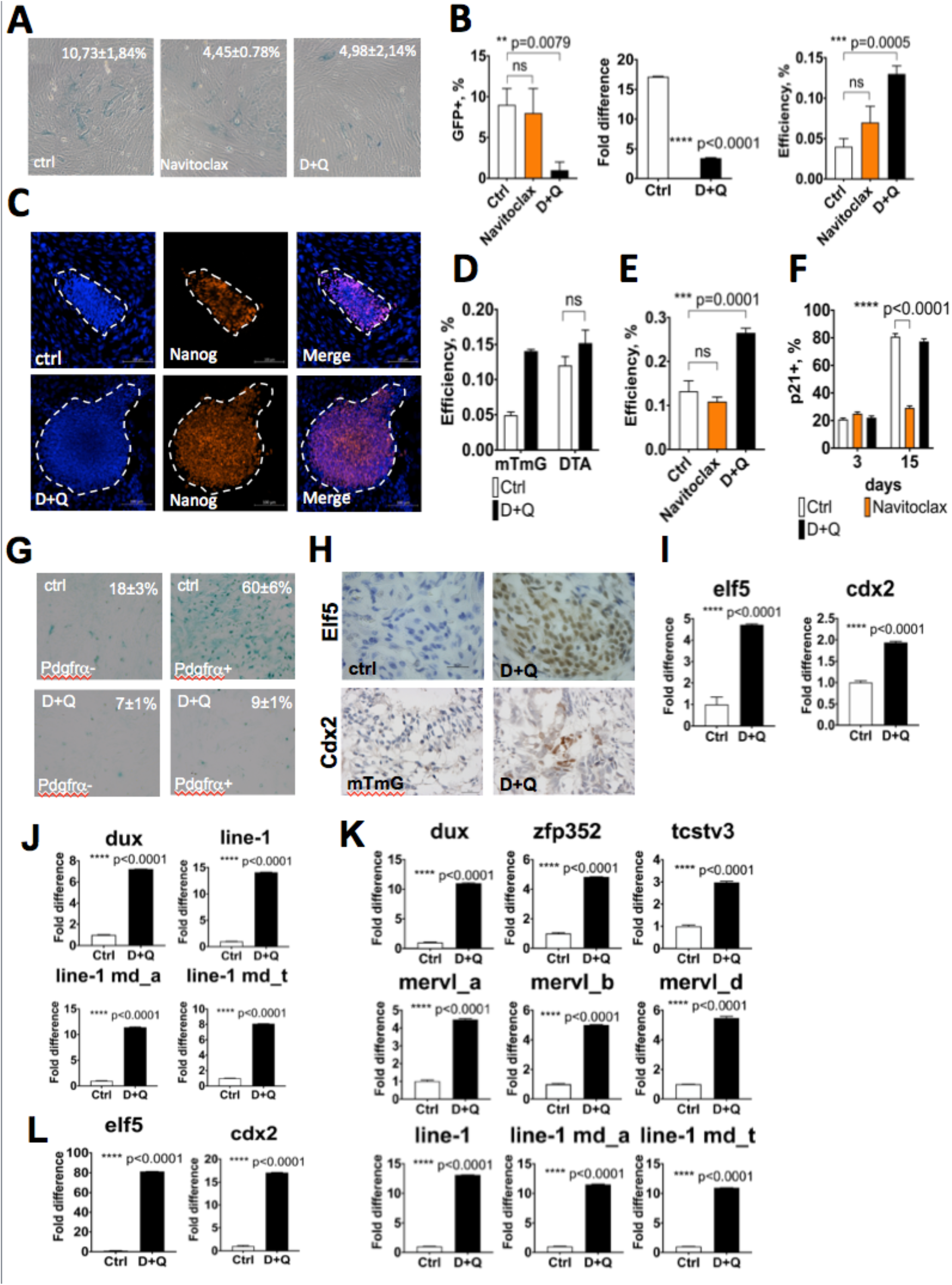
Chemical removal of p16^High^ cells during *in vitro* and *in vivo* reprogramming with senolytics dasatinib and quercetin (D+Q) leads to emergence of cells with totipotent features. **A** - analysis of SA-β-gal+ cells (± SD) in mock- and D+Q-treated DFs on d15. **B** (left panel) - analysis of accumulation of p16^High^ cells (± SD) in the presence of indicated drugs on d15. ns – not significant, p-value was calculated by ordinary one-way ANOVA with Sidak’s multiple comparisons test. Middle panel – qPCR analysis of p16 mRNA (± SD) in mock- and D+Q-treated DFs on d12. p-value evaluated by unpaired, two-tailed Student’s t-test with Welch’s correction. Right panel – efficiency of iPS reprogramming (± SD) of mock- and D+Q-treated DFs. ns – not significant, p-value was calculated by ordinary one-way ANOVA with Sidak’s multiple comparisons test. **C** - analysis of Nanog+ cells in iPS colonies derived from mock- and D+Q-treated DFs. Scale bar represents 100 μm. **D** - efficiency of iPS reprogramming of mTmG and DTA DFs with and without D+Q. Error bars represent SD, ns – not significant. **E** - efficiency of iPS reprogramming of p53-/-;rtTA;i4F DF, nock-treated (Ctrl) or treated with navitoclax or D+Q. ns – not significant, p-value was calculated by ordinary one-way ANOVA with Sidak’s multiple comparisons test. **F** - analysis of p21^High^ cells (±SD) during iPS reprogramming in DFs mock-, navitoclax-, or D+Q-treated. p-value was calculated by 2-way ANOVA with Sidak’s correction, **** p<0.0001. **G** - SA-β-gal staining of Pdgfrα+ and Pdgfrα– cells (±SD), isolated from teratomas developed in control mice or mice gavaged with D+Q. **H** – analysis of elf5 and cdx2 proteins in teratomas developed in control mice or mice gavaged with D+Q. Scale bars are 20 μm. **I** – qPCR analysis of elf5 and cdx2 mRNA in teratomas developed in control mice or mice gavaged with D+Q. Error bars represent SD, p-value evaluated by unpaired, two-tailed Student’s t-test with Welch’s correction. **J** – qPCR analysis of dux mRNA and non-LTR retroelement line-1 and its subclasses (±SD) in teratomas developed in control mice and mice gavaged with D+Q. p-value evaluated by unpaired, two-tailed Student’s t-test with Welch’s correction. **K** – qPCR analysis of 2C genes dux, zfp352, tcstv3 and the expression of LTR and non-LTR retroelements mervl and line-1 (±SD) in control and D+Q-treated iPS cells. p-value evaluated by unpaired, two-tailed Student’s t-test with Welch’s correction.

To confirm dependency of the D+Q effect on p16^High^ cells, we next reprogrammed DTA DFs with and without D+Q treatment. We found that treatment with D+Q had no additive effect on the efficiency of DTA iPS reprogramming, strongly implying that the role of D+Q in reprogramming is based on removal of p16^High^ cells (Figure 4D). It is well known that suppression of tumor suppressor pathways, including p53, is required to achieve successful reprogramming (Utikal et al., 2009). To confirm the dependency of D+Q on the presence of p53, we performed reprogramming of DF obtained from the p53-deficient pups expressing i4F. We observed further improvement of reprogramming of p53-deficient DF in the presence of D+Q, indicating that the mechanism of action of D+Q is p53-independent (Figure 4E).

The ability of navitoclax to reduce the number of SA-ß-gal^+^ cells without changing the number of EGFP^+^/p16^High^ suggested that it may target another senescence-related cell population that accumulates during iPS reprogramming. Recently, it was shown that navitoclax selectively reduced the number of p21^High^ but not p16^High^ cells in mouse liver while stimulating liver regeneration (Ritschka et al., 2020). Therefore, we evaluated the impact of navitoclax on p21^High^ cells during iPS reprogramming. Immunofluorescent analysis of p21 protein and EGFP (as a marker of p16^High^ cells) expression in DF revealed a significant increase in the number of p21-positive cells while <2% were p16/p21-double positive, suggesting that activation of p21 and p16 occurs in different cellular populations during iPS reprogramming. In turn, navitoclax caused a strong reduction in the number of p21^High^ cells during reprogramming (Figure 4F). Thus, our findings demonstrate that navitoclax efficiently removes p21^High^ cells, which appear to be non-essential in controlling the efficiency and quality of iPS reprogramming *in vitro*.

### D+Q clears p16^High^ cells in vivo and phenotypically mimics genetic removal of p16^High^ senescent cells during iPS reprogramming

Having shown that D+Q efficiently prevented emergence of p16^High^ cells during *in vitro* reprogramming in a manner similar to that achieved by genetic ablation with DTA, we next investigated the impact of D+Q *in vivo*. To assess the amount of EGFP^+^/p16^High^ cells present during *in vivo* reprogramming, mice were administered D+Q by oral gavage twice weekly immediately after DOX treatment until teratoma formation. Surprisingly, numerous mice died or had to be euthanized due to poor health while only few remaining mice developed teratomas. Analysis of teratomas revealed a significant reduction in the number of EGFP^+^/p16^High^ cells in D+Q-treated compared to vehicle-treated mice (SFigure 3A). Next, we extended our analysis of teratomas by isolating the Pdgfrα+ cell population. As shown in Figure 4G, Pdgfrα+ population purified from D+Q-treated mice showed a substantially less SA-ß-gal positivity compared to vehicle-treated mice, which was consistent with the reduced number of EGFP^+^/p16^High^ cells. Macrophage analysis revealed no difference between control and D+Q-treated teratomas (SFigure 3B) while H&E staining of D+Q-treated teratomas confirmed the presence of three germ cell layers, supporting the pluripotent status of reprogrammed cells (SFigure 3C). Further immunohistochemical analysis showed upregulation of the trophoblast stem cell marker Elf5 and Cdx2 in D+Q teratomas, and this was confirmed by qPCR analysis (Figure 4H&I). Similar to DTA teratomas, we also found a significant upregulation of 2C gene Dux and Line-1 mRNA in D+Q-treated teratomas (Figure 4J).

To characterize the properties of iPS cells after treatment with D+Q *in vitro*, we next reprogrammed DF in the presence of D+Q, and established feeder-free D+Q-treated iPS cell lines. We measured the expression levels of retro-elements as well as the 2C genes. As shown in Figure 4K, D+Q iPS cells showed increased expression of Mervl retro-elements, the 2C genes Dux, Zfp352, and Tcstv3 as well as Line-1 mRNA. We further evaluated the ability of D+Q iPS cells to differentiate into trophoblast stem cells and found significant upregulation of Elf5 and Cdx2 mRNA expression in D+Q iPS cells compare to control cells (Figure 4L). Thus, similar to DTA iPS cell lines, D+Q iPS cells exhibited several characteristics of totipotent-like cells with significant upregulation of expression of 2C genes, as well as LTR and non-LTR retro-elements.

To understand whether D+Q treatment induced the expression of the 2C genes and retro-elements in already established iPS lines, next we passaged (≥8) control iPS cells under naïve conditions in the presence of D+Q. Further analysis revealed no increase in the expression of the 2C genes, Mervl and Line-1 mRNA (SFigure 3D), suggesting that senolytic treatment can support the induction of a totipotent-like state only during reprogramming, but not in established iPS cells.

### Depletion of p16^High^ cells enhances widespread genomic H3K27 trimethylation and locus-specific H3K4 trimethylation during reprogramming

To gain insight into the molecular and genomic events that accompany reprogramming and that may be enhanced by depletion of p16^High^ cells (either genetically or by D+Q treatment), we analyzed changes in genome-wide H3K4 and H3K27 methylation at d12 after induction of reprogramming by addition of DOX. Using a spike-in chromatin-immunoprecipitation (ChIP) protocol to quantitatively compare global levels of each histone methylation, we observed that whereas total H3K4me3 levels are comparable to those of steady-state fibroblasts, cells undergoing reprogramming showed a marked and significant increase in total H3K27me3 levels (Figure 5A). This was consistent with the positive role of the polycomb protein Ezh2 in iPS reprogramming (Fragola et al., 2013). Notably, methylation of H3K27me3 was further augmented when reprogramming took place under conditions of depletion of p16^High^ cells, in line with the observed increase in reprogramming efficiency (Figures 2C&4B).

**Figure 5.**
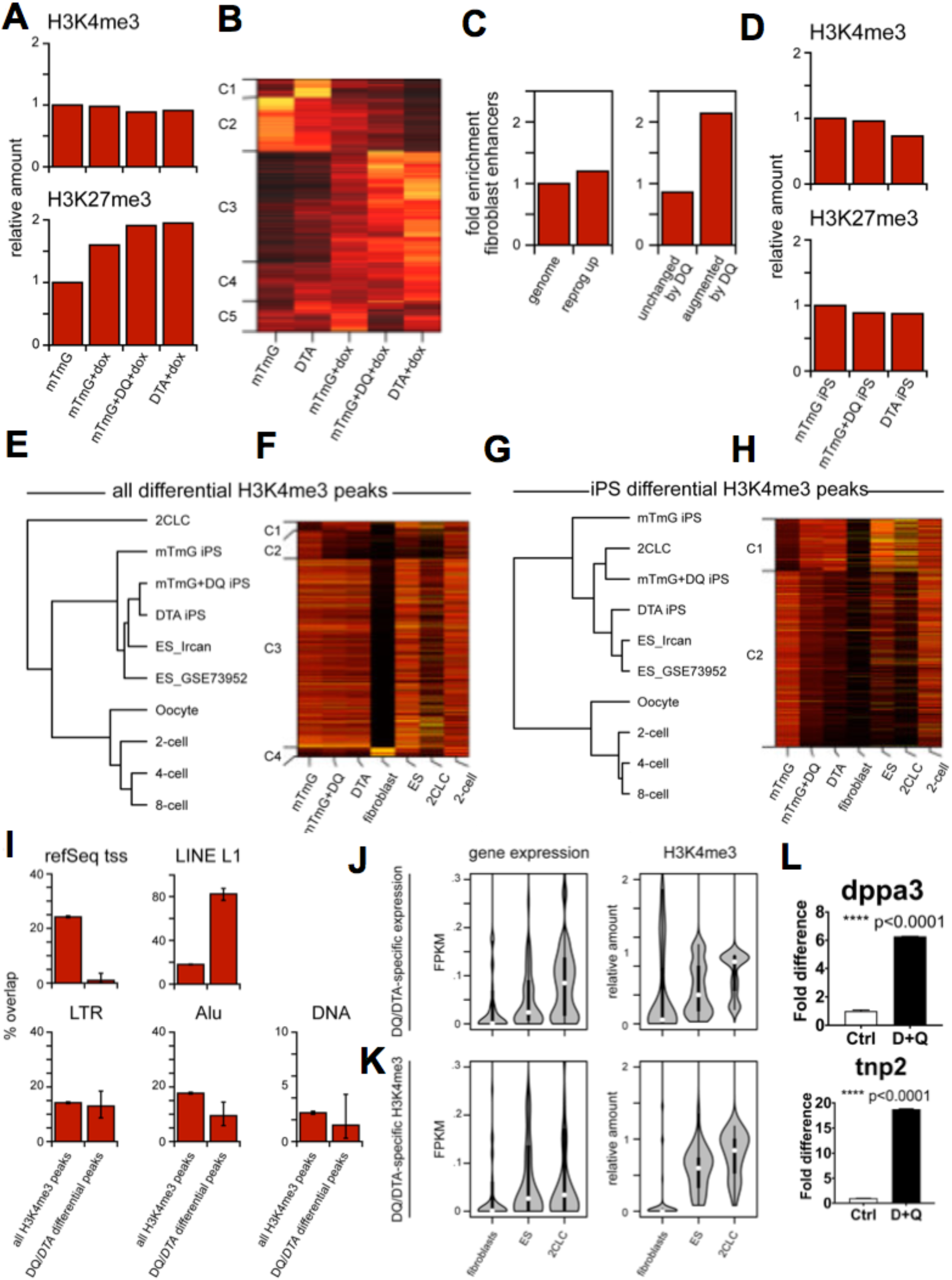
Depletion of senescent cells enables enhanced generation of iPS lines that share epigenetic characteristics of ES cells and 2-cell-like cells. **A** - quantitative measurement of global genomic H3K4me3 (top) and H3K27me3 (bottom) levels by spike-in ChIP-seq, in *mTmG* fibroblasts (‘mTmG’, left), *mTmG* fibroblasts at d12 of reprogramming induced by doxycycline treatment (‘mTmG+dox’, centre-left), *mTmG* fibroblasts at d12 of reprogramming, treated with dasatinib and quercetin (DQ; ‘mTmT+DQ+dox’, centre-right), and *DTA* fibroblasts at d12 of reprogramming (‘DTA+dox’, right). Increase in H3K27me3 associated with reprogramming (‘mTmG’ vs ‘mTmG+dox’): p<10^-300^; enhancement from DQ treatment (‘mTmG+DQ+dox’ vs ‘mTmG+dox’): p<10^-300^; enhancement in *DTA* cells (‘DTA+dox’ vs ‘mTmG+dox’): p<10^-300^ (Poisson test)). **B** - heatmap of relative H3K4me3 levels at peaks (rows) with differential coverage between samples (columns), with the 5 major clusters (C1-C5) of similarly-behaving peaks indicated. Note that the major clusters of peaks exhibiting decreases (C1, C2) or increases (C3,C4) in H3K4me3 during reprogramming display more-pronounced changes in experimental samples with depletion of senescent cells (‘mTmG+DQ+dox’ & ‘DTA+dox’). **C** - enrichment for enhancers linked to fibroblast-specific genes, in genomic loci that exhibit significant increases in H3K27me3 levels at d12 of reprogramming (left; corresponding to the top pie in panel g), and, among these, in loci that exhibit unchanged or augmented H3K27me3 increases upon DQ treatment (right; corresponding to blue & white pie slices in panel g). The expected enrichment based on genome-wide samping (‘genome’) is defined as 1. Augmentation of enrichment by DQ treatment: z-score 47.3 (by Monte-Carlo simulation), estimated p=1.3×10^-488^ (normal approximation); estimated p=1.03×10^-674^ (Poisson test assuming random distribution of enhancers). **D** - quantitative measurement of global genomic H3K4me3 (top) and H3K27me3 (bottom) levels by spike-in ChIP-seq, in iPS cells derived from *mTmG* fibroblasts (‘mTmG iPS’, left), *mTmG* fibroblasts treated with dasatinib and quercetin during reprogramming (DQ; ‘mTmT+DQ iPS’, centre), and *DTA* fibroblasts (‘DTA iPS, right). **E** - unsupervised hierarchical clustering of cell-types based on H3K27me3 levels at all differential peaks (relative standard deviation between samples ≥1; clustering based on Ward’s minimum variance). ‘ES_Ircan’ = ES cell data generated in this study; ‘ES_GSE73952’ and all embryonic cell-types = data from ncbi GEO series GSE73952, ‘2CLC’ = data from ncbi GEO series GSE166210, sample GSM5065951. **F** - heatmaps of relative H3K4me3 levels at all peaks (rows) with differential coverage between samples (columns). 4 major clusters (C1-C4) of similarly-behaving peaks are indicated. **G** - unsupervised hierarchical clustering of cell-types based on H3K27me3 levels at peaks with differential marking between iPS samples (see legend to panel c). **H** - heatmaps of relative H3K4me3 levels at peaks (rows) with differential coverage between iPS samples (first 3 columns: ‘mTmG’, ‘mTmT+DQ’ & ‘DTA’) – note that this corresponds to an enlarged view of clusters C1 & C2 in panel d. **I** - overlap of different classes of genomic elements with H3K4me3 peaks with increased H3K4me3 coverage in cells derived with depletion of senescent cells (corresponding to the most-different 200 peaks from cluster C1 in panel d). Reduced overlap with gene promoters (‘RefGene tss’): p<7.4×10^-124^; Elevated overlap with genomic instances of LINE-1-derived sequences (‘LINE L1’): p<3.1×10^-87^ (Binomial test). **J,K** - expression and H3K4me3 coverage in fibroblasts, ES-cells and 2C-like cells (‘2CLC’) at promoters of the top 50 genes that are specifically over-expressed (‘DQ/DTA-specific expression’, panel h) or that are specifically marked by elevated levels of promoter H3K4me3 (‘DQ/DTA-specific H3K4me3’, panel i) in iPS lines generated with depletion of senescent cells. Increased gene expression in 2CLC compared to ES cells: DQ/DTA-specific expression p<1.8×10^-2^; DQ/DTA-specific H3K4me3 p<6.7×10^-3^; elevated promoter H3K4me3 in 2CLC compared to ES cells: DQ/DTA-specific expression p=0.30; DQ/DTA-specific H3K4me3 p<4.4×10^-4^ (Mann-Whitney test). L-qPCR analysis of stella/dppa3 and tnp2 mRNA in control and D+Q-treated iPS cells. p-value evaluated by unpaired, two-tailed Student’s t-test with Welch’s correction.

Although global H3K4me3 levels were unchanged during reprogramming, the genomewide distribution of this mark changed significantly (Figure 5B, SFigure 4A). Hierarchical clustering confirmed a major change in H3K4me3 distribution between steady-state of both mTmG and DTA DFs and cells at d12 of reprogramming (SFigure 4B). Moreover, clustering based on H3K4me3 could robustly distinguish cells undergoing reprogramming with or without depletion of p16^High^ cells. H3K4me3 peaks themselves grouped into clusters with distinct patterns of behavior during reprogramming, and reprogramming-induced losses (Figure 5B; clusters C1,C2) or gains (clusters C3-C4) of H3K4me3 levels were consistently more-pronounced in conditions of depletion of p16^High^ cells. In particular, reprogramming-associated downregulation of H3K4me3 levels at the promoters of fibroblast-specific genes was significantly enhanced by D+Q treatment and in DTA cells (SFigure 4C). The H3K4me3 mark is a characteristic of transcriptionally-active promoters; thus, the presence of senescent cells counteracts reprogramming in part through impairment of genome-wide H3K4me3 redistribution.

In contrast to the highly locus-specific changes in H3K4me3 at peaks during reprogramming, we found that H3K27me3 levels exhibit widespread changes spanning broad multikilobase genomic domains (SFigure 4D). In line with the increase in total H3K27me3 levels at d12 of reprogramming (Figure 5A), up to 30% of all regions across the entire genome display reprogramming-induced increases in H3K27me3 (SFigure 4D&E). Regions gaining H3K27me3 predominantly correspond to loci with low H3K27me3 levels in fibroblasts under steady-state conditions (SFigure 4D, S4F): in contrast, regions with high levels of H3K27me3 before reprogramming are generally unchanged. Under conditions of depletion of p16^High^ cells this phenomenon was further augmented at a subset of the same set of genomic regions (SFigures 4D&E;4G). Furthermore, regions with augmented H3K27me3 levels upon D+Q treatment during reprogramming (or in DTA cells) displayed a highly-significant overlap with fibroblast-specific enhancers (Figure 5C) and other gene-regulatory elements. H3K27me3 is a mark linked to polycomb-associated gene repression, suggesting that widespread shut-off of fibroblast genes during reprogramming may be driven by broad H3K27me3 deposition, and that this is strongly enhanced in cell cultures lacking senescent cells. More generally, our data demonstrate that depletion of p16^High^ cells during reprogramming leads to major changes in histone methylation patterns, including H3K4me3 redistribution and H3K27me3 hypermethylation across a large fraction of the entire genome, and that this is likely a major contributor to the observed increased reprogramming efficiency.

### Depletion of senescent cells enables generation of iPS lines that share genomic and gene expression characteristics of both ES cells and 2CLC

The differentiation potential and marker expression of iPS lines that were generated under conditions of depletion of p16^High^ cells suggested that this setup may enable reprogramming towards a distinct cellular and genomic state. To investigate this, we used ChIP-seq and RNA-seq to interrogate the molecular phenotypes of iPS lines established under each condition, and to compare them to those of ES cell lines, of natural pre-implantation embryos, and of previously-reported transient- and chemically-induced 2-cell-like cell (2CLC) lines (Yang et al., 2022).

Global levels of H3K27me3 as well as H3K4me3 were equivalent in all iPS lines (Figure 5D), demonstrating that the augmentation in overall H3K27me3 levels observed at d12 of reprogramming after depletion of senescent cells (Figure 5A) is transient. However, both histone marks displayed notable differences in their genome-wide patterns according to the conditions of iPS generation (SFigure 4H). To address this in an unbiased fashion, we first used unsupervised hierarchical clustering of histone methylation levels at all differentially-marked peaks to assess the overall similarity of each iPS to cells derived from different pre-implantation stages of embryonic development, to ES cells, or to a transient 2CLCs. This revealed that, all iPS lines bear more similarity to ES cells than to any of the examined embryo cell-types or to 2CLCs (Figure 5E). Visualization of H3K4me3 levels at individual peaks revealed that while a large fraction of genomic loci acquire consistent marking resembling ES cells under all conditions of reprogramming, a subset of H3K4me3 peaks exhibit distinct behavior when reprogramming is performed under conditions of depletion of p16^High^ cells (Figure 5F; clusters C1,C2). We therefore repeated the hierarchical clustering of cell-types, but based only on the subset of peaks that are differentially marked between different conditions of iPS generation (Figure 5G). Using this approach, we found that the behavior of these peaks in D+Q-treated and DTA iPS lines resembles that of ES cells and 2CLCs, despite the overall genome-wide differences between iPS cells and 2CLCs. Examination of H3K4me3 levels at these peaks indicated that many peaks with very high levels of H3K4me3 in 2CLCs correspond to those that are preferentially marked in iPS cells generated under conditions of depletion of p16^High^ cells (Figure 5H), further implying that this subset of genomic loci may encompass elements that contribute to defining a totipotent-like phenotype.

Based on this, we analyzed the sets of genes and genomic features that are encompassed by H3K4me3 peaks that are differentially marked between control iPS cells and D+Q-treated/DTA iPS cells. Strikingly, we found that over 82% of the most-differential peaks correspond to genomic instances of Line-1 transposon-derived sequences (Figure 5I), compared to an average of only 18% at all H3K4me3 peaks in iPS cells. Other major classes of transposable elements, including LTR and Alu retrotransposons, and DNA transposons, were not enriched. The strong enrichment for H3K4me3 marking implies that deregulation of Line-1 loci at the chromatin level is a widespread phenomenon and represents a robust marker for this stage of cellular differentiation. In contrast, overlap of the most-differential H3K4me3 peaks with annotated gene promoters was significantly depleted (‘refGene tss’; Figure 5I), suggesting that the phenotype of iPS cells derived under conditions of depletion of p16^High^ cells may be driven by a small set of highly-specific genes.

We therefore focused on the sets of 50 genes defined by the highest level of upregulated mRNA expression (Figure 5J) or of elevated promoter H3K4me3 marking (Figure 5K) in D+Q-treated/DTA iPS cells. Remarkably, genes with elevated expression in D+Q/DTA iPS cells corresponded to those with significantly increased expression and H3K4me3 marking in totipotent 2CLCs, compared to ES cells (Figure 5J), consistent with a model whereby this set of genes plays a major role in specifying the totipotent potential of these cells. Genes with elevated H3K4me3 levels in D+Q-treated/DTA iPS cells (from cluster C1 in Figure 5H) displayed expression and H3K4me3 marking in both ES cells and 2CLCs, providing further support that these iPS cells share characteristics of both pluripotent and totipotent cell-types. Further analysis revealed 22 genes specifically enriched in D+Q-treated/DTA iPS when compared to ES and 2CLCs and several of them including a gene critical for early embryonic development, stella/dppa3 (Huang et al., 2017) were further verified by qPCR (Figure 5L, SFigure 4I).

Altogether, these genomic analyses define iPS cells established under conditions of depletion of p16^High^ cells as a distinct cellular state, which shares properties of both ES cells and 2CLCs, and which exhibit marking and expression of a highly-specific subset of genes that are likely to underpin their totipotent-like phenotype.

### Removal of senescent cells results in generation of iPS cells that contribute to all embryonic layers

Under defined chemical conditions, pluripotent stem cells with extended potency undergo formation of blastocyst-like structures(Sozen et al., 2019)(Sozen et al., 2018)(Li et al., 2019). These blastoids share some characteristics of 2.5 dpc mouse blastocysts, such as Nanog+ cell polarization, and appearance of Gata6+ and Cdx2+ cells. Since DTA and D+Q iPS cells are unique in their epigenetic makeup (Figure 5E-L), next we evaluated whether any of the available protocols would allow them to self-organize into blastoids. First, we differentiated different iPS toward trophoblast stem cells and evaluated their capacity to self-organize with corresponding non-differentiated iPS cells (Sozen et al., 2019)(Sozen et al., 2018). Our analysis however revealed no significant polarization of Nanog^+^ cells in aggregates formed *in vitro;* similar results were obtained when we used DTA and D+Q iPS without prior differentiation into trophoblast stem cells (data not shown). Next, we accessed a protocol based on selforganization of stem cell cultures in the presence of an inducer of trophoblast differentiation, Bmp4 (Li et al., 2019). As expected, all blastoids expressed Cdx2, with uniform distribution in the outer layer (Figure 6A), while removal of Bmp4 from the differentiation medium resulted in loss of Cdx2-positive cells (not shown). The primitive endoderm marker Gata6 was expressed in DTA and D+Q blastoids but totally lacking in the control. Nanog^+^ cells were found ubiquitously throughout WT blastoids, but in sharp contrast were polarized to one side in the majority of DTA and D+Q blastoids. This polarization represents a correct pattern of Nanog expression similar to that of mouse blastocysts. Thus, blastoids from DTA and D+Q iPS cells, but not control iPS cells, showed features of 2.5 dpc mouse blastocysts, recapitulating the key process of preimplantation development.

**Figure 6.**
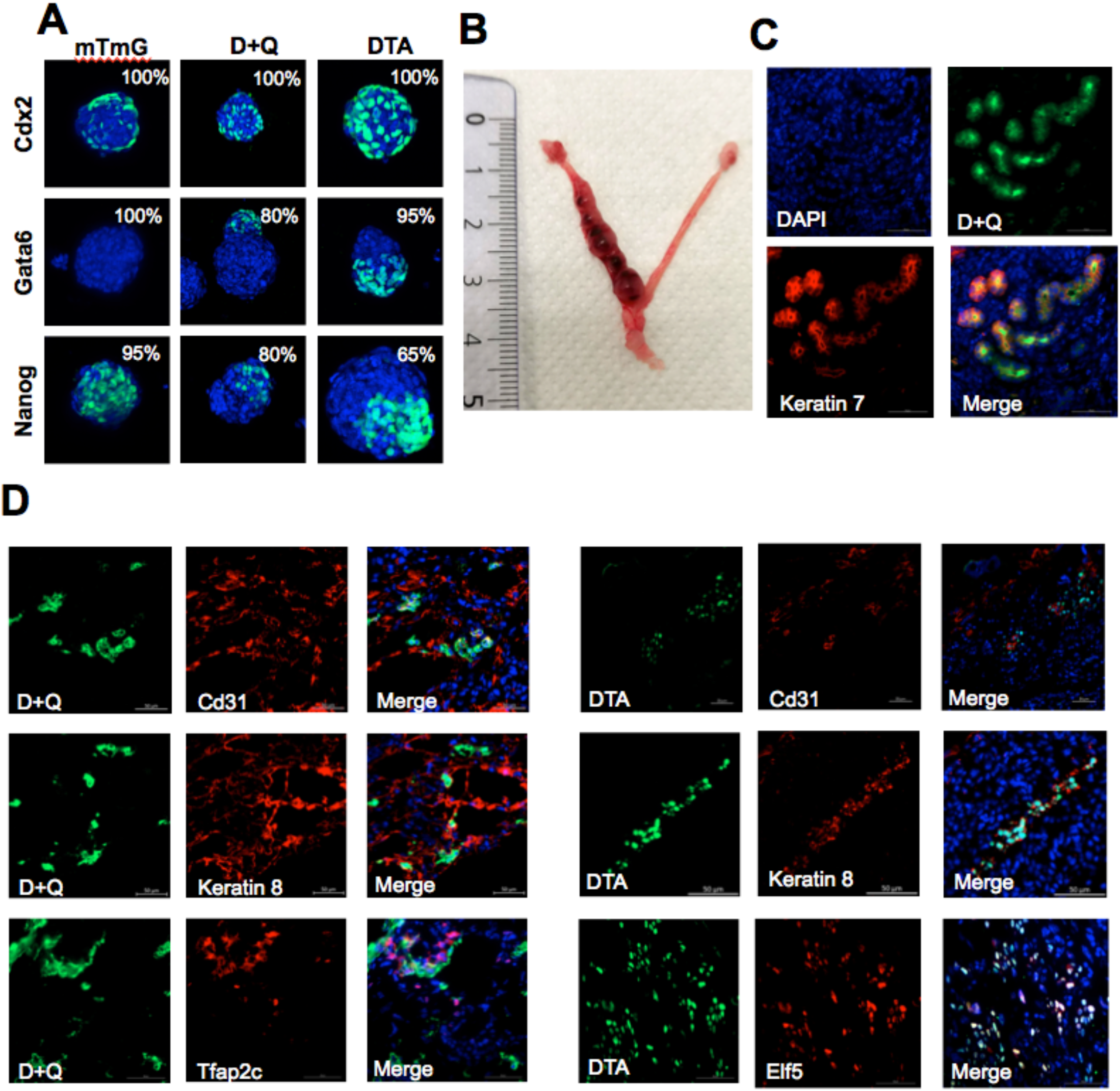
Senescence-free iPS lines produce implantation-competent blastoids *in vitro* and contribute to extraembryonic lineage *in vivo*. **A** - immunofluorescent analysis of blastoids obtained from mTmG, DTA and D+Q iPS stained for the trophoblast stem cell marker Cdx2, primitive endoderm marker Gata6 and pluripotent stem cell marker Nanog. Numbers represent percent of blastoids with shown pattern of marker expression. **B** – representative image of the uterus on d5 after implantation of GFP-expressing D+Q and DTA blastoids. **C** – immunofluorescent analysis of the implantation site of the uterus presented in B for Keratin 7, implantation marker, and GFP, marker of blastoid cells. Scale bar – 50 μm. **D** – immunofluorescent analysis of 12dpc placenta of chimeric embryos obtained by aggregation of GFP-expressing D+Q and DTA iPS with 8-cell morula from CD1 mice. Cd31 – vascular endothelium marker represents embryonic part of placenta. Keratin 8, Tfap2c, Elf5 – markers of an extraembryonic part of placenta. Scale bar – 50 μm.

Next, we investigated the ability of DTA and D+Q blastoids to undergo implantation and further embryonic development *in vivo*. We generated different iPS lines with stable expression of EGFP, then formed control, DTA and D+Q blastoids, which we implanted into the uterus of pseudo-pregnant mice. None of the control blastoids but around 10% of the DTA and D+Q blastoids implanted, induced decidualization and developed *in vivo* until at least 7.5 dpc (Figure 6B). Immunofluorescence analysis of the decidua as a hallmark of mammalian post-implantation embryo development confirmed the presence of EGFP^+^ cells at the implantation site identified by the expression of CK7 (Figure 6C). Further analysis, however, revealed no noticeable embryonic development beyond 7.5 dpc. At that stage, embryonic structures were surrounded by uterine tissue within the DTA and D+Q-blastoid-derived deciduae, suggesting that, DTA and D+Q blastoids exhibit limited developmental potential despite their ability to initiate implantation, which may be related to less-than-optimal culture condition of blastoids.

Recently, it has been proposed that more stringent conditions are required to assess the ability of totipotent-like cells to contribute to extraembryonic lineages(Posfai et al., 2021). To further confirm that DTA and D+Q-treated iPS cells possess an ability to contribute to development of extraembryonic tissues *in vivo, we* next used such stringent analysis after the aggregation of DTA and DQ iPS with diploid 8-cell morulas isolated from CD1 mice, and subsequent implantation into pseudo-pregnant females. As shown at Figure 6D, analysis of 12.5 dpc placentas revealed that senescence-free iPS cells contributed to both embryonic CD31-positive vascular endothelial cells, and trophoblast lineage cells labeled by a trophoblast progenitor marker Elf5 or pan-trophoblast markers CK8 and Tfap2c. Thus, our analysis confirmed that genetic or chemical removal of senescent cells results in generation of iPS cell lines with totipotent-like properties.

### Senescence-dependent regulation of NNMT during iPS cell reprogramming modulates S-adenosyl methionine (SAM) levels and totipotent-like properties

In the absence of p16^High^ senescent cells, the level of global methylation at K27me3 during reprogramming was significantly increased (Figure 5A-C). SAM is a universal donor of methyl groups, and its level decreases steadily with transition from the 2C to the late blastocyst state during early mouse embryonic development (Martinez-Val et al., 2021)(Zhao et al., 2021). Thus, we hypothesized that the effect of removal of p16^High^ senescent cells on reprogramming into a totipotent-like state could be linked to intracellular SAM levels. In testing this hypothesis, we found that SAM levels were significantly higher in DTA and D+Q-treated DFs compared to control DF on day 12 of reprogramming (Figure 7A). Similar increase in SAM levels was found in senescence-free teratomas and iPS lines (Figure 7B, SFigure 5A&B). It can be speculated that depletion of SAM by p16^High^ senescent cells could explain their negative impact on iPS cell reprogramming as this would cause a significant reduction in cellular plasticity due to their inability to provide sufficient methyl-donors for essential epigenetic and genetic methylation events. This, in turn, would be critical for efficient iPS cell reprogramming as well as for achieving totipotency as the highest level of stem cell potency. We next evaluated the consequences of blocking SAM synthesis using the Mat2 inhibitor, cycloleucine, on iPS cell reprogramming. As a second confirmation, we used a specific RNA-based SAM sensor (Dey et al., 2022), originally developed to detect intracellular levels of SAM, which appeared efficient in depleting intracellular SAM as well (SFigure 5C). We found that both cycloleucine and a SAM sensor significantly increased the number of p16^High^/EGFP-positive cells while blocking the efficiency of iPS reprogramming (Figures 7C&D; SFigure 5D&E)). Furthermore, cycloleucine was highly toxic during reprogramming of D+Q-treated DFs while depletion of SAM with an RNA sensor completely blocked any reprogramming. Only mTmG (control) and DTA DFs underwent iPS cell reprogramming in the presence of cycloleucine both with lower efficiency than nontreated DFs (Figure 7D, S5E). We further found that while cycloleucine treatment did not affect the level of expression of pluripotency genes in established iPS lines (SFigure 5F), it significantly reduced the expression of Line1 retro-elements and a 2C gene Dux in DTA iPS cells (Figure 7E). During blastoid formation, cycloleucine blocked the potential of Bmp4 to induce the expression of Cdx2 in WT iPS as well as fully inhibited the expression of Cdx2, Gata6 and polarization of Nanog+ cells in DTA iPS cells (Figure 7F). Thus, our data show that cycloleucine-mediated depletion of SAM levels reversed the ability of DTA DFs to be reprogrammed into totipotent-like DTA iPS cells.

**Figure 7.**
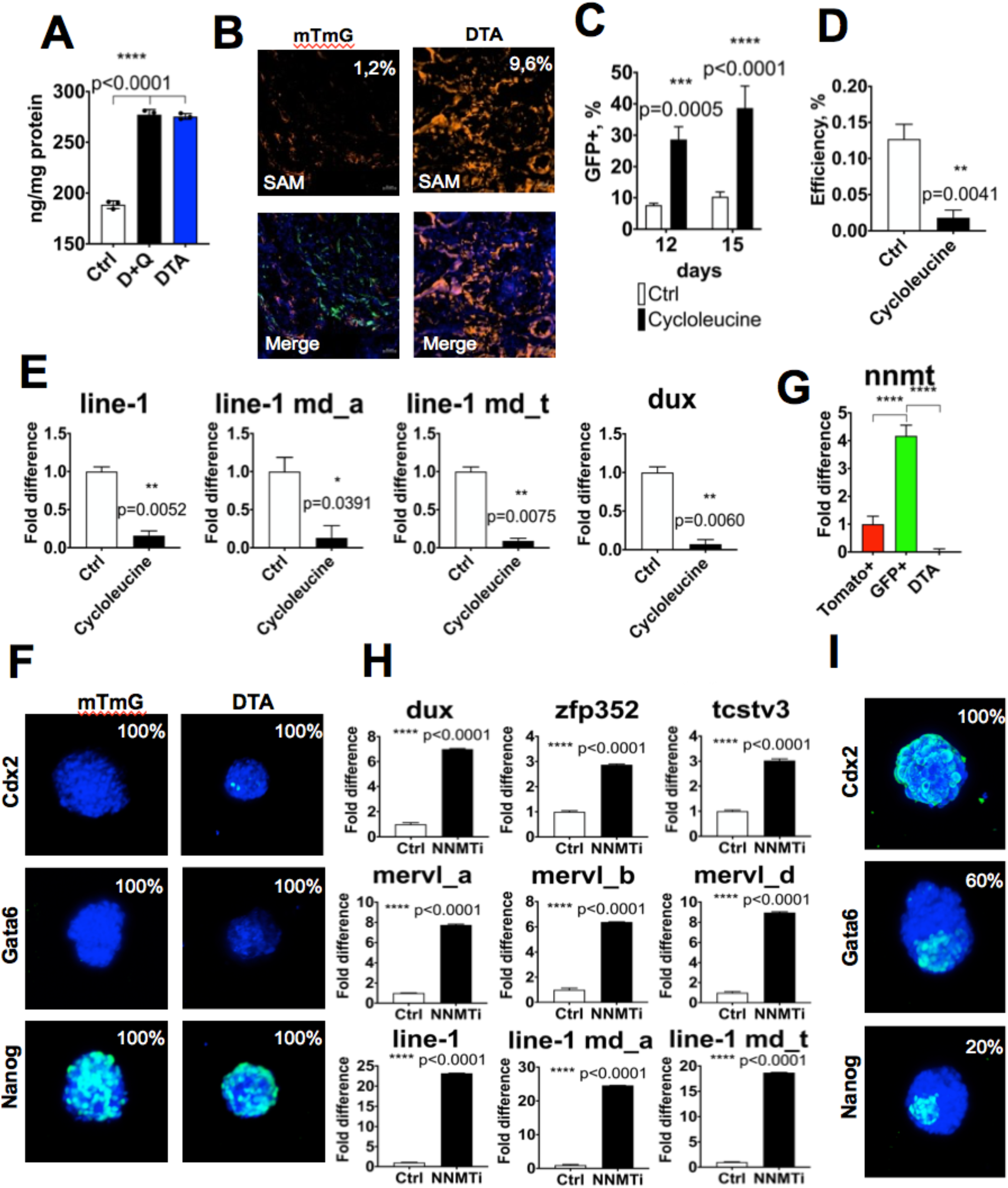
NNMT-dependent regulation of SAM blocks transition into a totipotent-like state during iPS reprogramming. **A** – quantification of SAM levels (±SD) by ELISA during iPS reprogramming in control, D+Q-treated and DTA DF on d12. p-value was calculated by 2-way ANOVA with Sidak’s correction. **B** – immunofluorescent analysis of SAM content in mTmG and DTA teratomas. Numbers - % of SAM-positive cells. **C** – analysis of p16^High^ cells (±SD) on d12 and d15 of reprogramming in the population of control and cycloleucine (5 mM) -treated DFs. p-value was calculated by 2-way ANOVA with Sidak’s correction. **D** – efficiency (±SD) of DTA DFs iPS reprogramming in the presence of 5 mM cycloleucine. p-value evaluated by unpaired, twotailed Student’s t-test with Welch’s correction. **E** – qPCR analysis of expression (±SD) of total, md_a and md_t subclasses of line-1 retroelement and 2C gene dux in control and cycloleucine (5 mM) –treated iPS. p-value evaluated by unpaired, two-tailed Student’s t-test with Welch’s correction. **F** - immunofluorescent analysis of blastoids obtained from mTmG and DTA iPS, derived in the presence of 5 mM cycloleucine. Numbers represent percent of blastoids with shown pattern of expression of analyzed markers. **G** – qPCR analysis of nnmt mRNA (± SD) in FACS-sorted GFP+ (p16^High^) and Tomato+ (p16^Low^) from p16Cre;i4F;rtTA;mTmG DFs and p16Cre;i4F;rtTA;DTA (DTA) DFs on d12. **** p<0.0001. P-values estimated by ordinary one-way ANOVA with Sidak’s multiple comparisons test. **H** – qPCR analysis of expression level of 2C genes dux, zfp352, tcstv3 and LTR and non-LTR retroelements mervl and line-1 in control and NNMTi-treated iPS cells. Error bars represent SD, p-value evaluated by unpaired, two-tailed Student’s t-test with Welch’s correction. **I** - immunofluorescent analysis of blastoids obtained from NNMTi iPS. Numbers represent percent of blastoids with shown pattern of expression of analyzed markers.

Next, we investigated the potential mechanism underlying the senescence-dependent regulation of SAM levels. After 12 days of reprogramming, we conducted FACS-sorting of mTmG DF and isolated Tomato+/p16^Low^ and EGFP^+^/p16^High^ populations for gene expression analysis. We found that NNMT mRNA, a SAM-consuming enzyme, was significantly upregulated in p16^High^ DF but considerably reduced in DTA DFs when compared even to p16^Low^ Tomato+ cells (Figure 7G). This implies that while NNMT upregulation occurs to a higher degree in p16^High^ senescent cells, these cells, through a paracrine mechanism, increase the level of NNMT in the rest of the cell culture. Indeed, multiple SASP factors that are produced by p16^High^ cells during reprogramming (Figure 2B) have been shown to upregulate NNMT expression (Roberti et al., 2021).

To investigate the role of NNMT further, we performed reprogramming of p16Cre;i4F;rtTA;mTmG DF in the presence of an NNMT inhibitor, JBSNF. The NNMTi enhanced the efficiency of iPS cell reprogramming (SFigure 5G), while established NNMTi-iPS cells contained high level of SAM and expressed increased levels of Mervl and Line-1 retro-elements as well as upregulated transcription of the 2C genes (Figure 7H). Furthermore, NNMTi-iPS cells formed blastoids *in vitro* with expression of Gata6-positive cells and polarization of Nanog^+^ cells (Figure 7I) similar to that observed in the DTA and D+Q iPS cell lines (Figure 6A). Thus, maintenance of high SAM levels during iPS cell reprogramming, which can be achieved by either depletion of senescent cells or inactivation of the SAM-consuming enzyme, NNMT, is sufficient for cellular reprogramming into a totipotent-like state.

## Discussion

To dissect the role of senescent cells in iPS reprogramming, here we employed genetic and chemical approaches to remove p16^High^ cells. In contrast to the suppressive effect of Cdkn2a deficiency, but consistent with a p53-deficient mouse background (Mosteiro et al., 2016), we found that the removal of p16^High^ senescent cells strongly accelerated 4F-induced reprograming *in vivo*. We further found that removal of p16^High^ cells resulted in an NNMT-dependent increase in SAM levels during the reprogramming, which in turn, allowed to achieve a high state of methylation-dependent epigenetic remodeling to support greater cellular plasticity. As a result, the iPS cell lines that emerged on senescence-deficient backgrounds expressed numerous markers of 2C-like cells with features of experimental totipotency.

Entry into an embryonic 2C-stage involves significant epigenetic changes and represents a critical *in vivo* developmental step in the transition from maternal to embryonic transcriptional control, a process known as zygotic genome activation (ZGA). One of the features of ZGA, and thus the totipotent 2C state, is developmentally programmed splicing failure that contributes to the attenuation of cellular responses to DNA damage, including reducing p53 pathway activation (Wyatt et al., 2022)(Shen et al., 2021). Similarly, activation of both p53- and p16-dependent checkpoints reduces the efficiency of iPS reprogramming (Li et al., 2009)(Banito et al., 2009)(Utikal et al., 2009) supporting the idea that cell cycle inhibitors represent important barriers to both iPS reprogramming and entry into the 2C-like state. Transient induction of inhibitory checkpoint molecules occurs in response to multiple stimuli, while sustained activation of both p53 and p16 is a feature of senescent cells that are broadly found during iPS reprogramming *in vitro* and *in vivo* (Li et al., 2009)(Banito et al., 2009)(Utikal et al., 2009)(Mosteiro et al., 2016).

A growing understanding of totipotency *in vivo* as well as 2C-like cells (2CLCs) which represent a small fraction (<1%) among ES and iPS cells when cultured under naïve conditions *in vitro* allowed to propose several approaches to move closer in establishing stem cell lines with totipotent potential (Genet and Torres-Padilla, 2020)(Riveiro and Brickman, 2020). Recent analysis of 2C embryos identified splicing failure as one of the features of the totipotent state and spliceosome inhibition using different approaches yielded the generation of stem cell lines with 2C-like potential(Shen et al., 2021). In addition, analysis of ES cells under naïve culture conditions *in vitro* revealed that several factors, such as a pioneer transcription factor, Dux, epigenetic factors Tet and Setdb1, or a chemical cocktail to produce expanded pluripotent conditions (EPSC) (reviewed in (Riveiro and Brickman, 2020)), all capable of regulating entry into a 2C-like state. Similar to 2CLC, the totipotent-like cells described here express high levels of pluripotent genes, with a transcriptome and a landscape of enriched histone H3K4me3 and H3K27me3 markers at regulatory elements that more closely resemble ES cells than 2C embryos (Figure 5)(Yang et al., 2022). However, there were clear difference between totipotent-like cells described here and 2CLCs, including the expression of Line1 retro-elements and Stella/Dppa3. Line-1 transcripts were found to be highly expressed and critical for regulation of global chromatin accessibility in the early mouse embryo (Jachowicz et al., 2017). In fact, the highest level of Line-1 transcripts was found in 2C embryos, with decreased levels by the 16C stage. In turn, the high Line-1 expression at early embryonic stages appeared to be essential for preimplantation development (Jachowicz et al., 2017)(Vitullo et al., 2012) and generation of somatic (Garcia-Perez et al., 2007), and germline mosaicism (Richardson et al., 2017) as a fundamental mechanism of genomic diversification. In a similar manner, Stella/Dppa3 is critical for early embryonic development, while its deletion results in significant attenuation of expression of MERVL in 2C embryos (Huang et al., 2017). Thus, in addition to upregulation of several 2C factors including Dux and MERVL typical of 2CLCs, totipotent-like cells described here express Line1 and Stella/Dppa3 to further re-enforce totipotent state.

It is noteworthy that the role of p16^High^ senescence in the regulation of totipotency found in this study does not reflect either the *in vivo* (during embryogenesis) or *in vitro* (shuttling in and out of 2CLC state in naïve ES cells) situations. Late embryogenesis has been shown to contain senescent cells, however they are not present during early embryonic stages (Muñoz-Espín et al., 2013)(Storer et al., 2013). Similarly, NNMT is expressed in the late stages of embryogenesis, while embryonic development is not impacted in NNMT knockout mice (https://www.mousephenotype.org/data/genes/MGI:1099443). Furthermore, we found no evidence of the presence of p16^High^ cells in established cultures of iPS lines maintained under different including naïve conditions. The role of p16^High^ senescence seems to converge with the regulation of totipotency at the level of SAM, a universal donor of methyl groups. Earlier studies showed the highest level of SAM in 2C embryos *in vivo*, which is consistent with an enhanced demand for epigenetic regulation at this embryonic stage including ZGA (Zhao et al., 2021). Furthermore, SAM levels seem to increase in the naïve 2CLC-containing state when compared to primed ES cells(Martinez-Val et al., 2021). We further found that senescence- and SAM-dependent regulation of pluripotency only exists during iPS reprogramming, while extended culturing of established iPS cell lines or ES cells in the presence of senolytic drugs dasatinib and quercetin, does not induce the appearance of totipotent properties (SFigure 3D). In this respect, the role of senescence in iPS cell reprogramming *in vitro* is somewhat different from that *in vivo*, since p16^High^ cells emerge before Nanog-positive cells *in vitro*, while the reverse situation is observed *in vivo*. This implies that *in vivo* reprogramming occurs without the induction of senescence and p16^High^ cells emerge only at the stage of teratoma formation; as such, *in vivo* reprogrammed Nanog-positive cells should be totipotent. Indeed, capturing early *in vivo* 4F reprogrammed cells from blood and subsequently culturing them *in vitro* has been shown to produce ES cells with totipotent potential (Abad et al., 2013). In contrast, when reprogrammed Nanog-positive cells start to produce teratomas *in vivo*, emerging p16^High^ senescent fibroblasts quickly derail the totipotent potential as the frequency of finding differentiated cells expressing extraembryonic markers Cdx2 and Elf5 is significantly lower or even absent in control mice when compared to mice without p16^High^ cells (Figures 3D&4H). It is also worth mentioning that depletion of SAM levels and lowering the SAM/SAH ratio via p16^High^ senescence-dependent and potentially independent mechanisms, could significantly attenuate cell plasticity as for example we observed that cycloleucine strongly reduced the ability of BMP4 to activate the expression of a trophoblast stem cell marker Cdx2 during blastoid differentiation (Figure 7F). Thus, high NNMT and low SAM availability not only restrict the potential of iPS cells to acquire totipotent properties, but also could attenuate cell fate transition during different physiological processes. In turn, targeting NNMT and SAM could go beyond current findings and have an impact on improving the plasticity and physiology of aged tissues and organisms.

## Acknowledgements

The authors acknowledge the IRCAN’s Molecular and Cellular Core Imaging (PICMI), Histology, Genomic and Animal Housing Facilities. The research in DB lab is supported by the Foundation FRM, Agence Nationale de la Recherche (grant “Senage”) and the INSERM program «Agemed». Research in SS lab is supported by the Agence Nationale de la Recherche. We are very grateful to Dr. Han Li (Pasteur Institute, Paris, France) and Dr.M.Serrano (IRB, Barcelona, Spain) for the i4F mouse strains (Abad et al., 2013).

## Declaration of interests

BB Grigorash and DV Bulavin filed a patent application related to this study

## Material and Methods

### Mouse models

The following lines of transgenic mice were used: p16Cre;mTmG and p16Cre;DTA (Grosse et al., 2020), i4F (strain B) (Abad et al., 2013), p53^-/-^ (Jackson Laboratory). Animals were kept in pathogen-free conditions during normal 12h light cycle with food provided ad libitum. For all experiments, mice heterozygous for all transgenes of both sexes no older than 6 months were used excluding for experiments where animal age was particularly specified. For allotransplanatation of iPS cells NSG immunocompromised mice were used. All animal experiments were performed in compliance with the Animal Care and Use Committee and approved by the ethical review committee.

### Derivation of primary mouse dermal fibroblasts

DF isolated from newborn (not older than 2 days after birth) pups were used as a starting culture to obtain iPS cells. Backskin of newborn pup was treated with dispase II (2.5 mg/ml in DMEM, Gibco) overnight at +4°C, after it was transferred to a collagenase IV solution (1 mg/ml in DMEM, Gibco) for 1 hour at room temperature. Collagenase was inactivated by addition of DMEM with 10% FBS and cell suspension was successively passed through nylon mesh strainers with pore diameter 100, 70, 40 μm. Cells were resuspended in DF culture medium (DMEM, 10% FBS, 1x penicillinstreptomycin, all from Gibco) and plated on 150 mm petri dish. This passage was considered zero andcells were frozen after genotyping.

### iPS reprogramming

DFs were seeded into 6-well plates (Falcon, Thermo Fisher) at 5×10^4^ cells per well in DF medium. Next day, medium was changed to iPS cell medium supplemented with doxycycline (1 μg/ml, Sigma-Aldrich). Medium for iPS cells contained: Knockout DMEM (Gibco # 10829018) supplemented with 15% ES-grade FBS (Gibco #116141079), 1x NEAA (Gibco #11140035), 0.055 mM beta-mercaptoethanol (Gibco #21985023), 1x penicillin and streptomycin solution (Gibco #15140122), 1x GlutaMAX (Gibco #35050061), 10 ng/ml mouse recombinant LIF (ES cell-tested) (Gibco #A35935). Upon reaching confluence, the cells were replated at the original density. Medium with doxycycline was changed every day. Senolytic drugs were used at following final concentrations: navitoclax, 0.1 μM (Selleckchem, #S1001); dasatinib, 100 nM (Selleckchem # S7782); quercetin, 1 μM (Selleckchem #S2347); JBSNF000088 (NNMT inhibitor), 1 μM (Selleckchem #S6779). All iPS cell lines were cultured in feeder-free conditions on Petri dishes covered with Matrigel (Corning, #354234) in media used for reprogramming with addition of 3 μM GSK3β inhibitor (CHIR99021, Selleckchem #S2924) and 1 μM MEK inhibitor (PD0325901, Selleckchem #S1036). Adaptation of D+Q iPS cell line to feeder-free conditions was performed in the presence of 10 μM ROCK inhibitor (Selleckchem #S8324).

### Trophoblast stem cell differentiation

10^4^ WT, DTA and D+Q iPS cells were plated per one well of Matrigel-coated (Corning, #354234) 6-well plate in trophoblast stem cell media (Tanaka et al., 1998) comprised of 70% mouse embryonic fibroblast-conditioned media (R&D #AR005) and 30% RPMI-1640 supplemented with 20% ES-grade FBS (Gibco #16141079), 1x Glutamax (Gibco #35050061), 1 mM sodium pyruvate (Gibco #11360070) and 0.055 mM b-mercaptoethanol (Gibco #21985023). 25 ng/mL FGF-4 (Peprotech, #100-31) and 1 μg/mL of heparin sulfate (Sigma, #H3149) were added to final media composition. Media was changed daily for 5 days, on last day cells were collected for analysis.

### Lentiviral particles packaging

Transfer vector along with packaging plasmids (psPax2, Addgene #12260 and p-CMV-VSVG, Addgene #8454) were contransfected into HEK 293T cells with FuGene HD transfection reagent (Promega #E2311). 24 hours after transfection, media was changed. Supernatant with viral particles was collected at 48 and 72h after transfection and cells were infected twice. Transduction efficiency was assessed by GFP expressing virus and subsequent quantification of GFP-positive cells by Countess II FL Cell Counter (Thermo Fisher).

### RNA isolation and qPCR

RNA was isolated using RNeasy Plus mini kit (Qiagen #74134) according to the manufacturer’s protocol. The RevertAid First Strand cDNA Synthesis Kit (Thermo Fisher #K1622) was used for cDNA synthesis. cDNA obtained during the reverse transcription reaction was diluted 5 times and 2-3 μl was used per 20 μl RT-PCR. RT-PCR was performed on the StepOne Plus Real-Time PCR System (Applied Biosystems) with reagents from KAPA SYBR FAST qPCR Kit (KAPA Biosystems #KK4618). Primer sequences used in the study are available upon request.

### Histological analysis

For paraffin embedding, tissue was fixed for 24 hours in 10% buffered neutral formalin, after it was transferred to 70% ethanol. Antigen epitope retrieval was performed in citrate buffer (10 mM sodium citrate, pH 6.0) or Tris-EDTA buffer (10mM Tris Base, 1 mM EDTA Solution, 0.05% Tween 20, pH 9.0). Endogenous peroxidase activity was blocked in 3% H_2_O_2_ solution in methanol for 15 min at RT. Sections of 4-5 μm thickness were blocked in 5% goat serum, permeabilized with 0.2% Triton X-100 and incubated with primary antibodies. Goat immunoglobulins against rabbit or mouse immunoglobulins conjugated with horseradish peroxidase (Promega) were used as secondary antibodies. Peroxidase activity was visualized using a DAB Substrate kit (Vector Laboratories). Nuclei were counterstained with Mayer’s hematoxylin (Sigma #MHS16). Slides were mounted using Pertex permanent mounting medium (Cellpath #SEA-0100-00A). Samples were analyzed with the Leica microscope in a brightfield mode. For embedding in OCT, samples were fixed in a 4% paraformaldehyde for 90 min on ice and subsequently transferred into 30% sucrose overnight at +4C. Next, tissue was snap-frozen in OCT (Thermo Fisher) and sectioned on a cryostat microtome (Thermo Fisher). For some antibodies epitope retrieving was performed. Blocking, permeabilization and incubation with primary antibodies was done the same as for paraffin sections. Secondary antibodies were used conjugated with a fluorescent label (Alexa Fluor 488, 546 or 647, 1:1000 Thermo Fisher). Nuclei were stained with DAPI. Slides were mounted with Fluoroshield mounting medium with DAPI (Sigma #F6057). Samples were analyzed using a Zeiss LSM880 confocal microscope or a Zeiss Axioskope fluorescent microscope. Cell cultures were fixed with ice-cold 4% paraformaldehyde for 5min, and immunofluorescent staining was performed similarly to cryosections. List of primary antibodies used: MacroH2a (Abcam #91528, 1:250), H3K9me2 (Abcam #1220, 1:250), γH2Ax (Cell Signaling #9718; 1:1000), Nanog (Cell Signaling #8822, 1:100), Desmin (Abcam #15200, 1:100), F4/80 (Abcam #210250, 1:400), GFP (Abcam #13970, 1:1000), aSMA (Abcam #124964, 1:100), Pdgfra (Miltenyi Biotec #130-101-905, 1:100), Ki67 (Abcam #16667, 1:50), Cd31 (Abcam #182981, 1:50), Gfap (Novus Bio #NB300-141, 1:1000), Elf5 (Elabsciense #E-AB-53122, 1:1000), Cdx2 (Abcam #76541; 1:300), Gata6 (Cell Signaling #5851, 1:500), Keratin 7 (Genetex #GTX110414; 1:250), Keratin 8 (DSHB Hybridoma Product TROMA-I; 1:500), Tfap2c (Santa Cruz #12762, 1:200), SAM (Cloud-Clone Corp #PAG414Ge01, 1:100).

### Alkaline phosphatase staining

Cells were washed with PBS briefly and fixed in paraformaldehyde-free fixative solution (Sigma-Aldrich #A5472) for 5 min at room temperature. Next, cells were incubated in premixed BCIP/NBT solution (Sigma #B6404) for 20 min at RT in the dark until dark brown staining revealed. Stained colonies were fixed with 4% paraformaldehyde and scanned on an Epson V500 scanner. Counting of alkaline phosphatase-positive colonies was performed using FiJi software for at least three independent experiments.

### Detection of SA-ß-gal activity

The cells were washed with PBS and fixed for 5 min in ice-cold 4% paraformaldehyde. Incubation in SA-ß-gal staining solution (40 mM sodium citrate pH=6.0, 5 mM K3[Fe(CN)6], 5 mM K4[Fe(CN)6], 2 mM MgCl2, 150 mM NaCl, 1 mg/ml x -gal) lasted from 24 to 48 hours, depending on the rate of staining development, at +37C. After stained cells were fixed for 5 min with ice-cold 4% paraformaldehyde, and cell nuclei were stained with DAPI. Representative pictures of the stainings were acquired on Evos digital microscope in fluorescent and brightfield modes. Analysis of the total number of cells (based on DAPI) and SA-ß-gal positive cells was carried out using FiJi software for three independent experiments.

### Isolation of Pdgfra-positive cell population

A teratoma was cut into smallest possible pieces and dissociated for 1 hour at +37C in a collagenase A solution (Roche, 0.1 mg/ml in DMEM). The resulting cell suspension was passed through mesh strainers with pore diameter from 100 μm to 40 μm. After last strainer, cell pellet was resuspended with 2% FBS in PBS. Next, cells were incubated for 1 hour on ice with magnetic beads coated with antibodies against Pdgfra (Miltenyi Biotec #130-101-547). Separation of Pdgfra-positive and negative populations was performed with LS column (Miltenyi Biotec #130-042-401) placed in a magnetic rack (MACS Multistand, Miltenyi Biotec # 130-108-933). The isolated populations were resuspended in fibroblast medium, plated and used for analysis of SA-ß-gal activity.

### FACS sorting

p16Cre;i4F;rtTA;mTmF DF were detached from the plastic with Tryple solution, resuspended in 2% FBS in PBS with addition of 1 mM EDTA and subjected to sorting for GFP-positive and GFP-negative populations with BD Facs Aria 3 cell sorter. Sorted GFP-positive and - negative populations were used for RNA isolation.

### Formation of blastoids

Protocol developed by Li (Li et al., 2019) was used as a guideline. 6 000 mTmG, D+Q, DTA or NNMTi iPS cells were resuspended to single cells and seeded into well of Aggrewell 400 plate (Stemcell Technologies #34415) in 1 mL of complete blastoid media: 50% Embryomax KSOM (Sigma #MR-106), 25% N2B27 basal media, 25% complete TSC media supplemented with 12.5 ng/mL FGF-4 (Peprotech #100-31), 5 ng/mL BMP-4 (Peprotech #120-05), 10 ng/mL VEGF-165 (R&D 293-VE-010), 1 μg/mL heparin (Sigma #H6279), 5 μM ROCK inhibitor (Selleckchem ##S8324), 3 μM GSK3b inhibitor (Selleckchem # S2924), 0.5 μM A83-01 (Axon Medchem #1421). N2B27 basal media consist of Knockout DMEM (Gibco # 10829018) mixed with Neurobasal (Gibco #21103049) in 1:1 ratio, supplemented with 1x N2 supplement (Gibco #17202001), 1x B27 supplement (Gibco #17404044), 1x NEAA (Gibco #11140035), 0.055 mM beta-mercaptoethanol (Gibco #21985023), 1x penicillin and streptomycin solution (Gibco #15140122), 1x GlutaMAX (Gibco #35050061). Complete TSC media consist of RPMI-1640 (Gibco #A1049101) supplemented with 20% ES-grade FBS (Gibco #116141079), 1x NEAA (Gibco #11140035), 0.055 mM beta-mercaptoethanol (Gibco #21985023), 1x penicillin and streptomycin solution (Gibco #15140122). After addition of cell suspension, plate was spinned down at 300 g for 1 minute. Next day media was replaced with 2 mL of complete blastoid media without ROCK inhibitor. 96h later blastoids were subjected to immunofluorescent staining or to embryo transfer.

### Transfer of blastoids into pseudo pregnant mouse

CD1 female mice were crossed with vasectomized CD1 males. 12h after mating females were checked for vaginal plugs. Mice with plugs were housed individually and used for the blastoid transfer at 2.5 dpc. Transfer was conducted via non-surgical approach using mNSET device (Paratechs #60010). Total amount of 40-50 individually selected blastoids were used for one transfer. Recipient mice were sacrificed 5 days after transfer (7.5 dpc) and uterus was examined for the signs of embryo implantation. Successfully implanted blastoids were fixed and embedded in paraffin for further histological analysis.

### iPS cells aggregation with diploid 8-cell morula and embryo transfer

Prior to mating, CD1 female mice were superovulated following standard protocol. 12h after mating with CD1 fertile males mice were checked for vaginal plugs and pregnant mice were separated. Next day embryos were flushed from the infundibulum zone of the uterus and prepared for the aggregation. Meanwhile, iPS cells were dissociated with StemPro Accutase (Gibco #A1110501) into clumps of small number of cells. 8-cell morulas without zona pellucida were put individually in the depressions of Petri dish together with iPS clumps in the drop of KSOM media under oil (Ovoil, Vitrolife #10029). 24h later incorporation of GFP-positive iPS cells into embryo were checked under fluorescent microscope. Double-sided (6 embryos per side) embryo transfer was performed into the oviduct of pseudo pregnant mice via surgical approach. Mice were sacrificed at 12.5 dpc, placentas were dissected and processed for the histological analysis.

### Analysis of SAM levels by ELISA

8×10^5^ HEK293 cells were transfected with 5 μg plasmids with Fusion6 transfection reagent or 5×10^6^ Dermal Fibroblasts of different genotype were used on day12 of reprogramming. In some cases, cycloleucine (20mM) was added for 24h. In other cases we used SAM sensor plasmids to lower the level of SAM (a kind gift from Samie R. Jaffrey (Department of Pharmacology, Weill Medical College, Cornell University, New York, NY, USA) (Dey et al., 2022). For HEK293 cells we used control (pAV-U6+27-Tornado-F30-Squash-Broccoli) or SAM sensor plasmids (pAV-U6+27-Tornado-F30-Broccoli-Squash-SAM sensor 4-2 or sensor 5-1) to selectively lower the level of SAM. For Dermal Fibroblasts, fragments from original riboswitch pAV plasmids, containing human U6 promoter, RNA-aptamer and stop for PolII signal were re-cloned into Sal1/Not1 sites of lentivirus pHAGE based backbone, containing Blasticidin resistance gene. Lentivirus was produced in HEK293T cells with pPax2 and pVSVG helper plasmids.

For SAM analysis, two days after transfection, the cells were collected in 1 ml cold PBS and lysed by sonication 3 cycles of 30sec with amplitude 30 on ice. Cleared cell lysate was stored in −80° until analysis. SAM concentration in 50 μl cell lysate was analysed by S-Adenosylmethionine (SAM) and S-Adenosylhomocysteine (SAH) ELISA Combo Kit (Cell Biolabs) according to manufacturer protocol.

### Spike-in Chromatin immunoprecipitation (ChIP), sequencing, and data processing

Cells were washed twice with PBS at room temperature, and fixed by addition of formaldehyde to a final concentration of 1% to cells in 5 ml PBS and incubation for 15 min at room temperature. Fixation was stopped by addition of glycine to a final concentration of 125 mM. Fixed cells were washed once quickly and twice for 10 min each with ice-cold PBS, and collected by scraping and centrifugation for 10 min at 500g at 4 °C. Cells were resuspended at approximately 1 × 10^7^ ml^-1^ and nuclei released by incubation in 1 ml of ice-cold L1 buffer (50 mM Tris pH8, 2 mM EDTA, 0.1% NP40, 10% glycerol + protease inhibitors) for 5 min. Nuclei were collected by centrifugation for 5 min at 500g at 4 °C and resuspended at 5 × 10^7^ ml^-1^ in 900 μl of ice-cold L2 buffer (50 mM Tris pH8, 5 mM EDTA, 1% SDS + protease inhibitors). Chromatin was fragmented by sonication to an average size of 600–700 bp (typically 9 cycles of 10 s sonication, 1 min recovery on ice, using a micro-tip sonicator) and insoluble debris was pelleted by centrifugation. A 50 μl aliquot was removed from each sample and analysed by agarose gel electrophoresis after DNA extraction to verify fragmentation. Fragmented chromatin was diluted with 9 volumes of buffer DB (50 mM Tris pH8, 200 mM NaCl, 5 mM EDTA, 0.5% NP40) and pre-cleared using 40 μl of protein-A sepharose for 1 h at 4 °C with rotation. Each chromatin sample was mixed with a ‘spike-in’ of 10% (by initial cell number) chromatin prepared in an identical fashion from human HEK-293 cells. After centrifugation at 500g for 1 min, 4 μg of antibody was added to each 1 ml of supernatant containing the pre-cleared samples and incubated overnight at 4 °C with rotation. Antibodies used for ChIP were: Abcam 8580 (for H3K4me3), Upstate 07-449 (for H3K27me3). Antibody-bound chromatin was pulled-down by addition of 15 μl of protein-A or -G sepharose for 30 min at 4 °C, and collected by centrifugation. Chromatin-bound beads were washed once quickly and 4 times for 5 min each with 1 ml of ice-cold buffer WB (20 mM Tris pH8, 500 mM NaCl, 2 mM EDTA, 1% NP40, 0.1% SDS), followed by 3 washes for 5 min each with 1 ml of ice-cold TE. Immunoprecipitated chromatin was released by incubating beads in buffer EB (TE + 2% SDS) for 5 min at room temperature with periodic tickling, and the supernatant was collected. This was repeated two more times and the supernatantw were pooled. Fixation was reverted by overnight incubation at 65 °C, and DNA was directly purified using the MinElute PCR purification kit (Qiagen), and eluted in 30 μl elution buffer. DNA samples were quantified using picogreen reagent and a Qubit fluorometer (Invitrogen). ChIP efficiency was verified by qPCR with primers specific for known positive and negative control regions according to the antibody target. Sequencing libraries were prepared using the NebNext Ultra II DNA library kit (New England Biolabs), and sequenced using a paired-end strategy on a NextSeq500 instrument (Illumina).

Demultiplexed sequence data was aligned separately to the mouse mm9 and human hg19 genomes using bowtie (with options -v 2 -a -m 5 --maxins 2000 --tryhard); alignment rates were typically around 80%. The ratio of mouse:human aligned reads in input samples was used to determine the exact level of spike-in for each sample, and then the ratio of mouse:human aligned reads in the ChIP samples was used to calculate the relative overall level of ChIP recovery in each sample (Figures 5A,D). Excess duplicate reads that mapped to the exact same genomic location (corresponding to likely PCR duplicates) were eliminated if there were significantly more than expected based on the local read density in a range of ±2 bp at a p value of 0.05; the level of redundant reads that were filtered-out at this stage was typically around 10%. Genome-wide tracks of coverage were normalized by the sequencing depth and by the overall level of ChIP recovery in each sample (Figures S4A,C,E). Tracks of H3K27me3 coverage-differences (Figure S4C) were calculated as the pairwise differences in normalized coverages between samples in sliding windows with a step size of 1kb and covering a ±15kb interval.

### RNA sequencing and data processing

Cells were lysed directly into RNA lysis buffer (38% phenol, 0.8 M guanidine thiocyanate, 0.4 M ammonium thiocyanate, 0.1 M Na acetate pH5, 5% glycerol). Total RNA was separated by addition of 0.2 volumes of chloroform, emulsification by shaking, and centrifugation at 13,000g for 10 min at 4 °C, followed by precipitation of RNA in the upper, aqueous phase by addition of 1 μl of glycogen azure (Sigma) plus 2.5 volumes of isopropanol, centrifugation at 13,000g for 20 min at 4 °C, and washing the RNA pellet in 70% ethanol. RNA samples were dissolved in 20 μl water, and quantified using a Nanodrop spectrophotometer (Thermo). Strand-specific, poly-A enriched sequencing libraries were prepared using the TruSeq Stranded mRNA Library Prep kit (Illumina), and sequenced using a paired-end strategy on a NextSeq500 instrument (Illumina).

Demultiplexed sequence data was aligned to the mouse mm9 genome using star; alignment rates were typically around 77%. Aligned fragments mapping to refGene annotated transcripts were quantified using featurecounts (with options -M -O –fraction) and used to calculate fragments per kilobase of transcript per million aligned reads (FPKM) for each annotated transcript.

### Genomic data analysis

#### Peak calling

H3K4me3 ChIP-seq peaks were identified from deduplicated aligned reads separately for each sample using macs1.4 (with options p 1e-4 –nomodel –shiftsize=250 –keep-dup=all). Peaks from multiple samples were merged using mergeBed (bedTools), adjusted to a consistent width of 1kb centred on the peak summit, and annotated with the normalized ChIP coverage for each sample.

#### Selection of differential peaks

Peaks with differential coverage between samples (Figures 5B,E-H, S4B) were defined based on the relative standard deviation (rsd; sd divided by mean) of the coverages for all samples being considered. Cut-off rsds were selected based on the distribution for each set of data, and correspond to 0.5 (for H3K4me3 peaks during reprogramming; figures 5B,S4B), 0.8 (for H3K27me3 levels in all iPS, ES & 2CLC samples; figure 5E,F) and 1.0 (for H3K27me3 levels only in iPS samples; figure 5G,H).

#### Hierarchical clustering

Clustering was performed using the R hclust function with default parameters (euclidean distance, Ward’s method for clustering).

#### Defining specific sets of enhancers, promoters, genomic intervals and peaks

Fibroblast enhancers (Figure 5C) were defined as H3K4me1 peaks in unstimulated 3T3 fibroblasts (ncbi GEO dataset GSM801534). Fibroblast-specific promoters (figure S4X) were defined as the annotated start positions ±500bp of refSeq transcripts with FPKM >1 in 3T3 fibroblasts and FPKM <0.01 in ES cells. Genomic intervals that exhibit reprogramming-induced or senescence-affected changes in H3K27me3 levels (Figure S4D) were defined as 1kb bins with a mean difference in coverage between samples exceeding a cutoff >1000 aligned, normalized bp per 10M aligned reads. DQ/DTA-differential H3K4me3 peaks (Figure 5I) were defined as the 200 peaks with the highest ratio of mean H3K4me3 coverage between the mean of DQ & DTA iPS samples and control (mTmG) iPS cells. The enrichment of L1-derived sequences and the depletion of refSeq tss show in figure 5I is robust across a wide range of ratio cut-offs. DQ/DTA-specific promoters were defined based on the expression levels of the associated transcript (figure 5J) or on their coverage by H3K4me3 (Figure 5K), in each case selecting the 50 promoters with the highest ratio between DQ & DTA iPS samples and control (mTmG) iPS cells.

